# Diverse infection models demonstrate robust resistance of *Mycobacterium tuberculosis* to innate immunity

**DOI:** 10.1101/2025.11.20.687493

**Authors:** Marian R. Fairgrieve, Ella C. Brydon, Roberto A. Chavez, Dmitri I. Kotov, Russell E. Vance

## Abstract

*Mycobacterium tuberculosis* (Mtb) is a robust activator of innate immunity. However, there is little evidence that innate immune mechanisms control Mtb before the onset of adaptive immunity. Prior work has generally used specific pathogen-free (SPF) mouse models and relatively large infectious doses, which may obscure the capacity of innate immunity to control Mtb. Here, we performed ultra-low dose Mtb infections and found that the initial innate immune response was unable to curb even minimal Mtb infectious doses. Additionally, we primed the immune systems of C57BL/6 mice by co-housing with “pet shop” mice prior to Mtb exposure. Co-housed mice were as susceptible to Mtb infection as SPF mice. To pre-activate innate immunity at the site of Mtb infection more specifically, we also infected the lungs of mice with *Legionella pneumophila* (Lp) prior to Mtb. Innate immunity alone can clear large doses (>100,000 CFU) of Lp from the lung within a few days. However, priming the mouse lung by pre-infection with Lp only modestly reduced Mtb CFU compared to mice infected with only Mtb, indicating that Mtb can robustly replicate even in the presence of a strong anti-bacterial innate response. We performed single-cell RNA-sequencing on myeloid cells from mice either infected with Mtb alone or mice primed with Lp. We found that Lp priming before Mtb infection induced measurable changes in myeloid cells responding to Mtb, but these changes had little effect on innate control of Mtb. Together, these data demonstrate the robust resistance of Mtb to innate immune clearance under diverse experimental conditions.

## Introduction

Mtb robustly engages and activates innate immune cells. Diverse pattern-recognition receptors (PRRs) including Toll-like receptors, inflammasomes, NOD1/2, cGAS and dectin receptors all detect and respond to Mtb infection [1]. Engagement of PRRs by Mtb results in a robust inflammatory response. *In vivo*, Mtb initially infects and replicates within alveolar macrophages (AMs), which are thought to provide a permissive niche for Mtb replication in the initial days post-infection, while simultaneously inducing pro-in-flammatory responses [2-4]. Pro-inflammatory response can be further enhanced by contained Mtb (co-Mtb) or BCG vaccination infection models [5-7]. However, even in naïve, co-Mtb, or BCG-vaccinated mice, Mtb replicates many thousand-fold prior to the onset of adaptive immune control, which occurs starting after day 14. Genetic evidence also supports the conclusion that Mtb is largely resistant to early innate immune responses. Mice with genetic deficiencies in the innate immune response have nearly identical CFU burdens to wild type mice before the onset of adaptive immunity [8-17]. The existing data therefore suggest that although Mtb elicits a robust innate inflammatory response, Mtb replication is unrestricted during the early innate immune response.

The failure of innate immunity to control Mtb replication in the lung is not due to an intrinsic inability of the lung to control bacterial infections. Indeed, even large doses (>10^5^ CFU) of other pathogenic bacteria, including *Pseudomonas aeruginosa* and *Staphylococcus aureus* are readily and rapidly cleared from the lungs of mice [18, 19]. *Legionella pneumophila* (Lp) provides a particularly interesting example of a bacterial pathogen that is readily cleared by innate immune responses in the lung. Like Mtb, Lp initially infects alveolar macrophages and engages a very similar set of host pattern recognition receptors (the LPS of Lp does not activate TLR4) [20-22]. In contrast to Mtb, which establishes chronic infections even after an initial inoculum of 1-3 CFU, doses as high as 10^6^ CFU of Lp are cleared from the lungs of SPF C57BL/6 mice within 3-5 days post-infection [23, 24]. Many of the key innate pathways known to be critical for Lp clearance, such as TNFα, IL-1, and IFNγ, are also elicited by Mtb and help control Mtb infection. However, these pathways only aid in control during the adaptive phase of the response instead of during the innate phase [25-27].

Most studies of Mtb infection in mice have been performed with specific pathogen-free (SPF) C57BL/6 mice, using doses 20-100 times the infectious dose 50 (ID50), which is as low as 1 CFU in mice [24, 28]. Ultra-low dose (ULD) Mtb infections have been used as a promising model for evaluating BCG vaccine efficacy in mice, suggesting that the ULD infection model could be used as a sensitive method to uncover *in vivo* mechanisms of Mtb control [29]. In addition, laboratory mice kept in clean SPF facilities have underdeveloped immune systems compared to wild, or pet shop raised mice [30]. When C57BL/6 mice are co-housed with pet shop mice, their immune systems mature to more closely resemble the activation state of the immune system of adult humans, with increased numbers of circulating monocytes, macrophages, neutrophils, and memory T cells [30, 31]. Co-housing SPF mice with pet shop mice can strongly enhance protection against certain intracellular bacterial infections, though this effect is variable [32].

In this study, we systematically investigated whether the use of more physiologically relevant animal models could reveal a role for innate immunity in control of Mtb replication. First, we used an ULD infection model to test whether high infectious doses used in previous studies had obscured a potential contribution of innate immunity to Mtb control [24]. However, we found no evidence that innate immunity controls Mtb even at ultra-low infectious doses. As an alternative approach to enhance innate responses, we primed the innate immune system through either pet shop mouse co-housing or Lp co-infection. Although both models enhanced anti-bacterial innate immune responses, we found no evidence that a primed innate immune system impaired Mtb replication during the innate phase of the infection. Together, our results indicate that Mtb robustly resists innate immune clearance mechanisms in the lung, even under experimental conditions where the innate immune system is highly activated, such as during ultra-low dose infections or conditions in which lung immune responses are already primed.

## Results

### Even at ultra-low doses, innate immunity does not control Mtb

We hypothesized that non-physiological infectious doses of Mtb used in conventional mouse infections may have overwhelmed the mouse innate immune system, thereby obscuring the contribution of innate immunity to Mtb control. To sensitively test of the ability of innate immunity to control Mtb, we used a previously described method to infect wild-type C57BL/6J mice with ultra-low doses (1-3 CFU) of Mtb (Erdman strain) [29]. To assess whether innate immune responses restrict Mtb at these ultra-low doses, in parallel we also infected *Ifngr1*^*–/–*^, *Tnfr*^*–/–*^, *Sp140*^*–/–*^, *MyD88*^*–/–*^ *Trif*^*–/–*^, and *Atg5*^*fl/fl*^ *LysM-Cre*^*+*^ mice. These mice lack key factors involved in innate immunity, but prior work with conventional doses had only previously documented their importance in Mtb infection during the adaptive phase [33-37]. We reasoned that use of ultra-low doses may reveal a phenotype for these knockout mice during the innate phase (i.e., by day 14). In all cases, however, the immune-deficient mice had comparable CFU to control wild-type mice at 14 days post-ultra-low dose infection (Fig 1A-E). In addition, the fraction of mice that were successfully infected (i.e., harboring detectable CFU on day 14) were not significantly different when comparing wild type to immune-deficient animals (Table 1). In these experiments, 21%-62% of the mice remained uninfected across experiments (Table 1), consistent with an ultra-low or near ultra-low infectious dose (1-3 CFU) [24, 38]. Thus, the innate immune response to Mtb appeared to be remarkably ineffective *in vivo* and failed to control infections seeded by 1-3 bacteria.

**Table 1.**
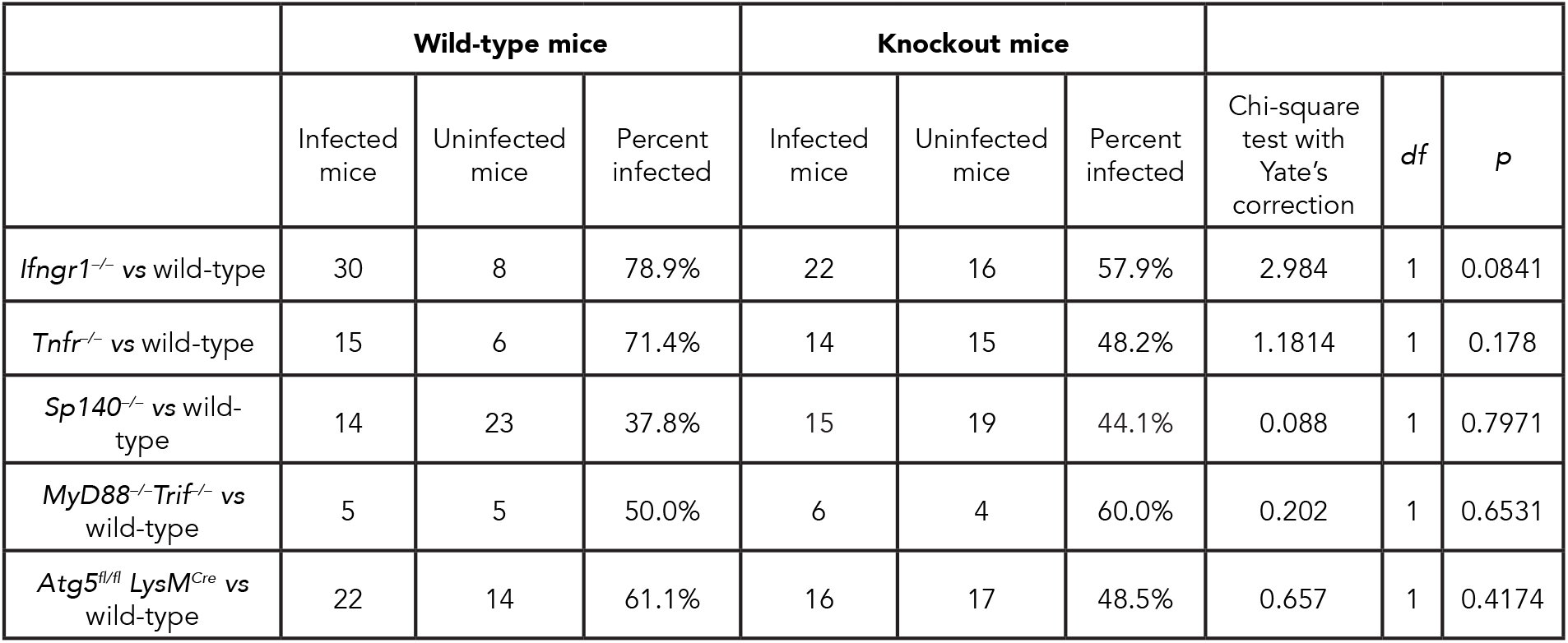
Calculation of the fraction of infected mice after ULD aerosol infection. Percent of infected wild-type or knockout mice and the results of chi-square tests with Yate’s correction of CFU positive and CFU negative wild-type or innate immune deficient mice infected with an ultra-low dose of Mtb. A 2 × 2 contingency table with used to calculate the results of the chi-square test.

|  | Wild-type mice |  |  | Knockout mice |  |  |  |  |  |
| --- | --- | --- | --- | --- | --- | --- | --- | --- | --- |
|  | Infected mice | Uninfected mice | Percent infected | Infected mice | Uninfected mice | Percent infected | Chi-square test with Yate's correction | df | p |
| *Ifngr1<sup>-/-</sup>* vs wild-type | 30 | 8 | 78.9% | 22 | 16 | 57.9% | 2.984 | 1 | 0.0841 |
| *Tnfr<sup>-/-</sup>* vs wild-type | 15 | 6 | 71.4% | 14 | 15 | 48.2% | 1.1814 | 1 | 0.178 |
| *Sp140<sup>-/-</sup>* vs wild-type | 14 | 23 | 37.8% | 15 | 19 | 44.1% | 0.088 | 1 | 0.7971 |
| *MyD88<sup>-/-</sup>Trif<sup>-/-</sup>* vs wild-type | 5 | 5 | 50.0% | 6 | 4 | 60.0% | 0.202 | 1 | 0.6531 |
| *Atg5<sup>fl/fl</sup> LysM<sup>Cre</sup>* vs wild-type | 22 | 14 | 61.1% | 16 | 17 | 48.5% | 0.657 | 1 | 0.4174 |

**Fig 1.**
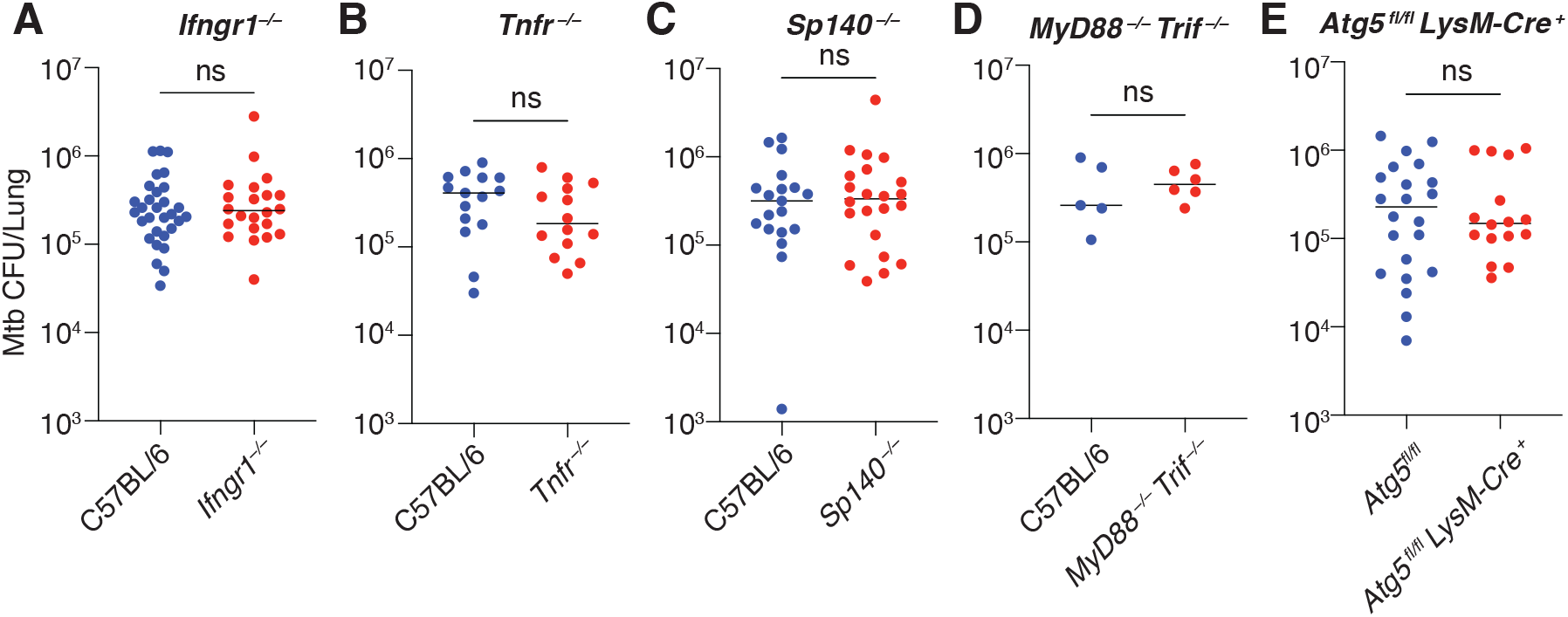
Innate immune knockout mice are not more susceptible to Mtb than wildtype. (**A-E**) Colony forming units (CFUs) of Mtb in the lungs of wildtype C57BL/6 and (A) *Ifngr1*^*–/–*^ (B) *Tnfr*^*–/–*^ (C) *Sp140*^*–/–*^ (D) *MyD88*^*–/–*^*Trif*^*-–/–*^ or (E) *Atg5*^*fl/fl*^ and *Atg5*^*fl/fl*^ *LysM-Cre*^*+*^. Lungs were analyzed 14 days post infection. The bars represent the median where CFU was detected. Pooled data from one to three independent experiments are shown. A Mann-Whitney test was used to calculate significance. *p ≤ 0.05.

### Priming C57BL/6 mice with pet shop mouse co-housing does not alter control of Mtb

We sought to test the hypothesis that innate immunity does not contribute significantly to initial Mtb control (up to day 14 post-infection) in SPF mice because the innate immune compartment is insufficiently matured or primed prior to Mtb infection. To mature the immune system of SPF mice, we employed a previously described pet shop mouse co-housing model [30]. We purchased female mice from a local pet shop in Berkeley, CA, and co-housed them with female SPF B6 mice for 60 days. Serological surveys for common mouse pathogens were performed, and pet shop mice indicated the presence of numerous pathogens including helicobacter, mouse parvovirus, murine norovirus, and rotavirus. The presence of pathogens was heterogenous across individual mice (STable 1). Co-housed mice displayed symptoms of pathogen exposure, including weight loss, particularly in the first 10 days of co-housing (SFig 1).

We measured the impact of co-housing by examining the number of innate and adaptive cells in the lung by flow cytometry after 60 days of co-housing (Fig 2A). Co-housing did not have profound effects on the innate immune cellular compartment in the lung at day 60 (Fig 2B). However, at this timepoint, there were significant differences in the number of CD4^+^ and CD8^+^ effector memory cells (Fig 2C), similar to the changes observed in previously published studies of pet shop mice, suggesting that the innate immune system has been activated beyond baseline [30, 32]. These changes varied by cage. Formerly SPF mice closely resembled the pet shop mouse with which they were co-housed (SFig 2A-D).

**Fig 2.**
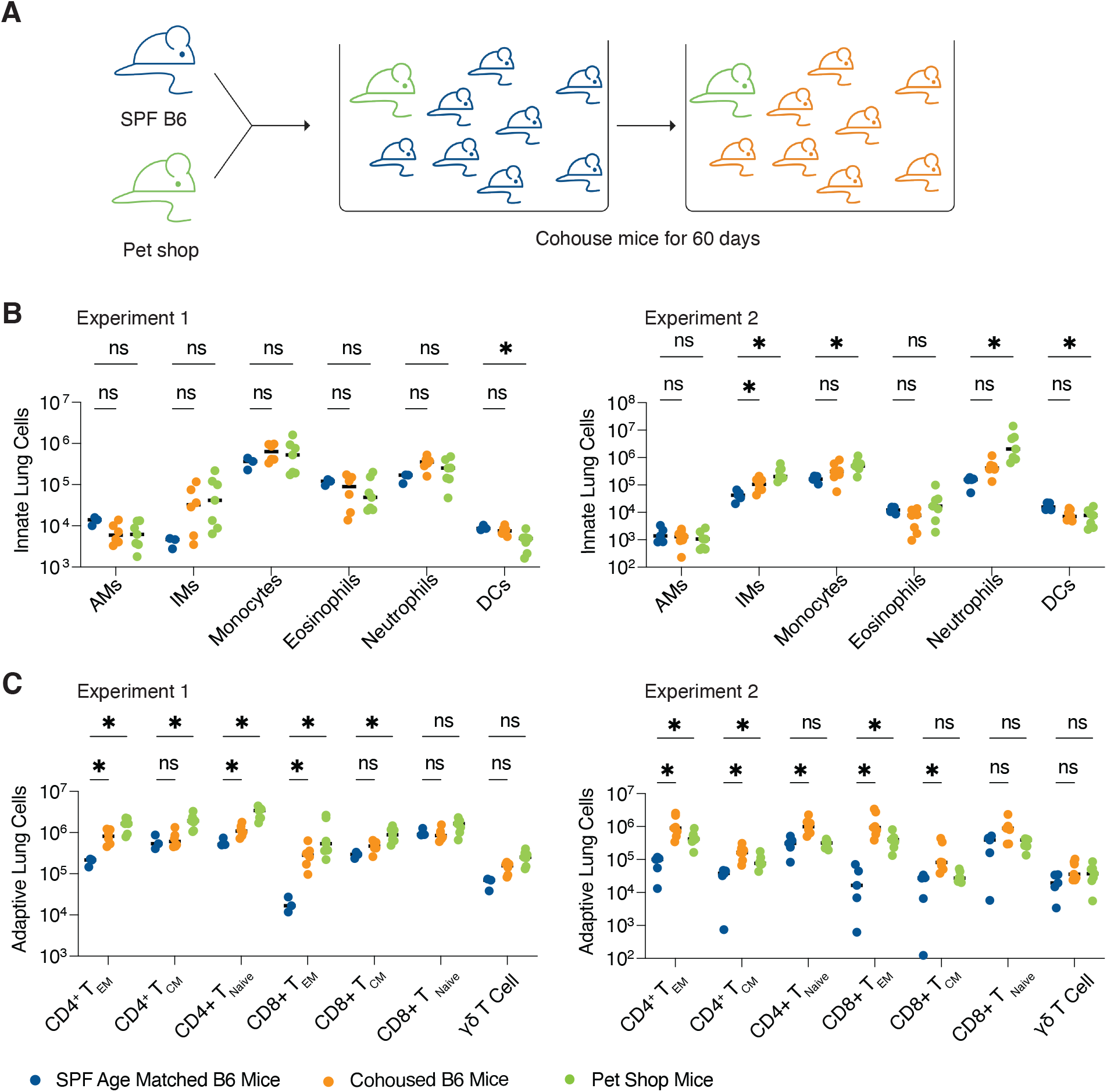
C57BL/6 mouse immune system can be primed by cohousing with pet shop mice. (**A**) Schematic of cohousing set up. Female C57BL/6 mice were cohoused with adult female pet shop mice for 60 days, during which co-housed mice are exposed to pathogenic, commensal, and opportunistic microbes carried by pet shop mice. (**B-C**) Lung innate (B) and adaptive (C) immune cells were profiled from age-matched SPF C57BL/6 mice, cohoused C57BL/6 mice, or pet shop mice in two independent experiments. A One-way ANOVA on log_10_-transformed data, followed by pairwise comparisons using Tukey’s test was performed to determine significance. EM= Effector memory, CM = Central memory, *p ≤ 0.05.

To confirm that our pet shop mouse co-housing procedure could enhance innate immune control of bacterial infections [31, 32], we infected age-matched SPF and co-housed mice with *Listeria monocytogenes* (Lm). In both the liver (Fig 3A) and spleen (Fig 3B), co-housed mice exhibited significantly lower CFU at 48 hours post-infection compared to SPF mice. We also infected mice intranasally with Lp. Co-housed mice largely contained similar numbers of CFUs as SPF mice, with one exception. Of the five cages of co-housed mice infected with Lp across two independent experiments, mice from four cages had similar levels of Lp as SPF mice and mice from one cage harbored significantly lower CFU in the lung, suggesting that in some instances, priming of the innate immune system by co-housing can enhance clearance of Lp (Fig 3C, SFig 2E).

**Fig 3.**
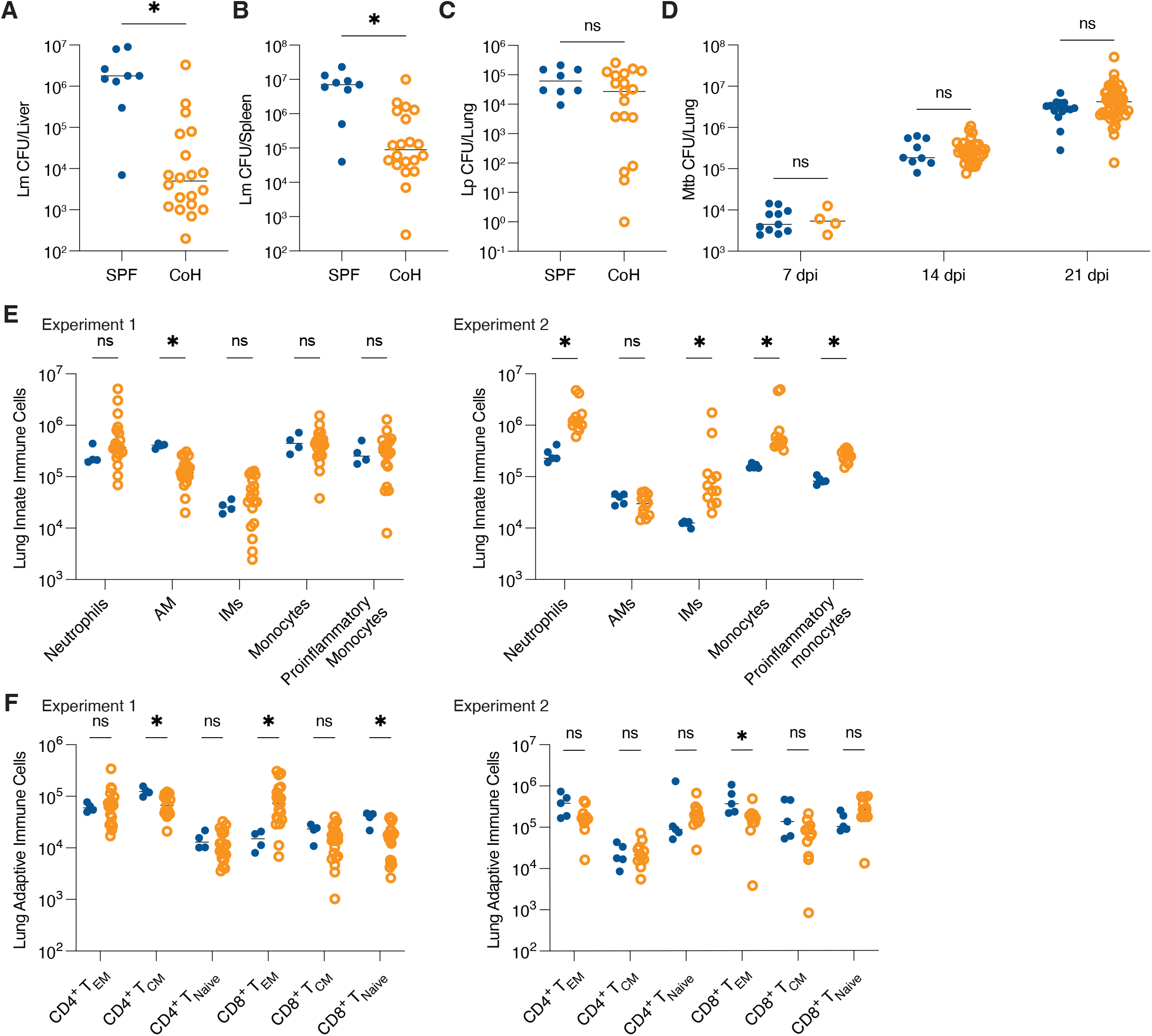
Pet shop cohoused mice exhibit enhanced control of Lm but not Mtb. (**A-B**) *Listeria monocytogenes* (Lm) CFUs from the liver (A) and spleen (B) from either co-housed (CoH) C57BL/6 mice or age-matched SPF controls. (**C**) Lp CFUs from the lung of either co-housed C57BL/6 mice or age-matched SPF controls. (**D**) Mtb CFUs from the lung of either co-housed C57BL/6 mice or age-matched SPF controls. (**E**) Quantification of innate lung cell populations from co-housed and SPF age matched mice from two independent cohousing experiments. (**F**) Quantification of adaptive lung cell populations from co-housed and SPF age matched mice from two independent co-housing experiments. The bars represent the median values. Multiple Mann-Whitney tests were used to calculate significance. *p ≤ 0.05.

We then infected a cohort of pet shop co-housed mice and matched SPF controls with Mtb. Because the number of surviving mice co-housed with each pet shop mouse was limited, we employed a standard low dose of 20-50 CFU to ensure all mice in each group were infected, and measured lung CFU at 7-, 14-, and 21-days post-infection. We found that co-housed and SPF mice contained similar numbers of CFU in the lungs (Fig 3D). Strikingly, the Mtb infections were very consistent, and CFUs did not vary substantially by cage, unlike mice infected with Lm or Lp.

We assessed the numbers of innate and adaptive immune cells in the lungs at 14 days post-Mtb infection (Fig 3E-F). The cellular response to infection in co-housed mice were variable compared to SPF mice, reflecting the variation seen in uninfected co-housed mice (Fig 2B-C), although differences cell numbers was generally reduced after Mtb infection. Together, our findings imply that co-housing with pet shop mice induces an altered immune state in SPF mice that can variably promote innate immune control of infections, but that Mtb is resistant to this shift.

### Mtb exhibits robust replication despite priming the innate immune system with *L. pneumophila*

Pet shop mice carry an array of viruses, bacteria, and parasites. The priming effects of these diverse pathogens may vary substantially or even counteract each other. Some may promote anti-bacterial immunity in the lung, whereas others may hinder it. In addition, each pet shop mouse may carry or transmit a distinct combination of pathogens, and it is therefore difficult to conduct controlled experiments. Moreover, although pet shop co-housing reduced systemic Lm colonization and increased the number of activated lymphocytes in the lung, there were inconsistent effects on Lp clearance in the lung. Therefore, it was unclear whether pet shop co-housing consistently induced an anti-bacterial innate immune response in the lung.

We therefore turned to co-infections with Lp, a pathogen that is established to elicit a robust and highly effective anti-bacterial innate immune response in the lung (SFig 3A-C). Co-infections with Lp avoided the complexities of co-housing and allowed us to activate innate immune responses without eliciting confounding antigen-specific anti-Mtb adaptive immune responses. Lp also infects and is cleared from the same cell type (alveolar and interstitial macrophages) that Mtb infects during the initial innate phase of infection (SFig 3C). We infected mice with Lp strain JR32, a clinically derived isolate of Lp [39]. Lp secretes flagellin into the cytosol of infected host cells, resulting in activation of the NAIP– NLRC4 inflammasome and induction of rapid pyroptotic cell death [40, 41]. To avoid killing alveolar macrophages, we infected mice with Lp lacking *flaA* (LpΔ*flaA*, hereafter referred to as Lp), which does not activate NAIP–NLRC4-mediated pyroptotic cell death [40, 41].

To confirm that priming with flagellin-deficient Lp enhances clearance of subsequent bacterial infections, we infected mice with Lp, followed by a second Lp infection 2 days later (Fig 4A).

**Fig 4.**
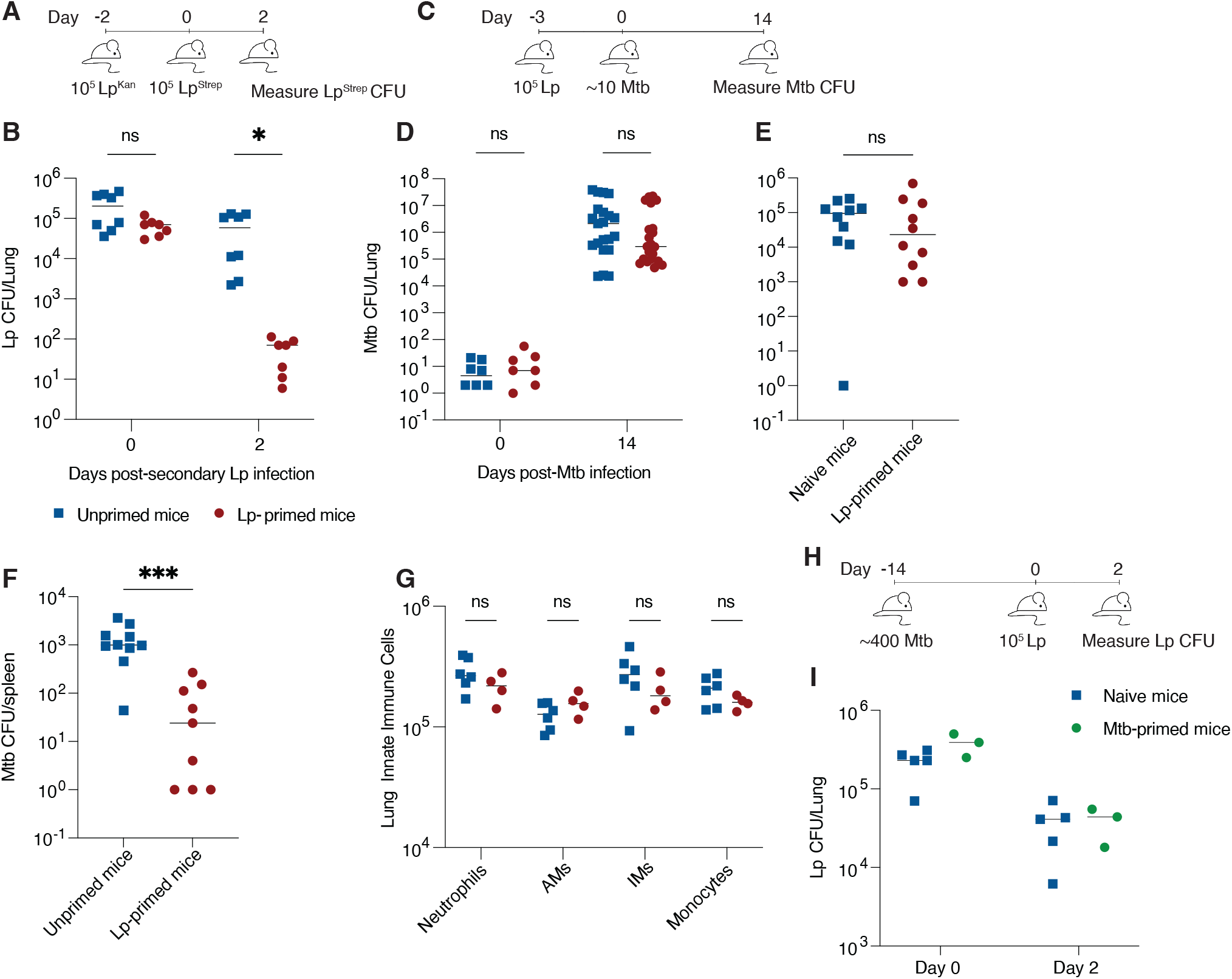
Priming with Lp does not curb Mtb replication. (**A**) Schematic of Lp co-infection. Mice were infected with Lp containing a kanamycin resistance gene (Lp^Kan^) or were left uninfected. Two days later, all mice were infected with Lp containing a streptomycin resistance gene (Lp^Strep^). Two days later, Lp^Strep^ CFUs were measured. (**B**) Lp lung CFU from unprimed mice or mice primed with a previous infection of Lp. Results shown are from 2 combined experiments. (**C**) Schematic of Lp/Mtb co-infection. Mice were infected with Lp or were left uninfected. Three days later, all mice were infected with Mtb. 14 days later, Mtb CFU was measured in the lung and spleen and samples taken for flow cytometry analysis. (**D**) Lung Mtb CFU from unprimed or mice primed with a previous infection of Lp. Results shown are from 3 combined experiments. (**E**) Lung Mtb CFU from unprimed or mice primed with a previous infection of Lp and infection with Mtb via the intranasal route. Results are shown from 2 combined experiments. (**F**) Spleen Mtb CFU from unprimed or mice primed with a previous infection of Lp and following aerosol infection of Mtb. (**G**) Lung innate immune cells from unprimed mice or Lp-primed mice. (**H**) Schematic of Mtb/Lp co-infection. Mice were infected with ∼400 CFU of Mtb or were left uninfected. 14 days later, all mice were infected with Lp. (**I**) Lp lung CFU from unprimed mice or mice primed with a previous infection of Mtb. The bars represent the median where CFU was detected. Mann-Whitney tests were used to calculate significance. *p ≤ 0.05.

Each infection was performed with a strain bearing a unique antibiotic resistance marker so the strains could be distinguished. While previous infection with Lp does not reduce the initial CFU that reach the lung during a secondary infection, we found that priming with Lp results in approximately 3-log lower CFU in the lungs 2 days post-secondary Lp infection compared with un-primed mice (Fig 4B). This implies that priming with Lp strongly activates an anti-bacterial innate immune response.

We then tested whether priming the lung with Lp could enhance control of Mtb compared to unprimed mice (Fig 4C). We first infected mice intranasally with 10^5^ CFU of Lp. At 3 days post-Lp infection, we then infected mice with approximately 10 CFU of Mtb via the aerosol route. By this time point, Lp has been cleared from the lungs by an inflammatory innate immune response (SFig 3A) [26, 42]. At both 1 and 14 days post-Mtb infection, we compared Mtb CFU between unprimed mice, which were infected only with Mtb, and Lp-primed mice. Strikingly, priming did not impact Mtb burdens in the lung at 14 days post-infection (Fig 4D), suggesting that the anti-bacterial lung environment induced by prior Lp infection did not inhibit Mtb replication. To determine if the route of infection could alter the effect of Lp priming, we primed mice with Lp via the intranasal route and followed with an intranasal Mtb infection 3 days later. Again, there was no difference in Mtb CFU in primed mice as compared to unprimed mice (Fig 4E).

Strikingly, while Lp priming did not affect Mtb CFU in the lung, it did reduce Mtb dissemination to the spleen at 14 days post-infection, suggesting Lp priming induced an immune response that partially restricted extra-pulmonary dissemination or replication (Fig 4F).

We measured innate immune cells in the lungs by flow cytometry (Fig 4G). After 14 days of Mtb infection, there were similar numbers of neutrophils, interstitial macrophages (IMs), monocytes, and alveolar macrophages (AMs), in Lp-primed mice compared to unprimed mice. Only small increases in cytokine expression were detected in the lungs of both unprimed and Lp-primed mice as compared to uninfected mice at the day 14 post-infection. Large increases in key cytokines induced by Mtb infection were only detected at later time points (SFig 4). As a complementary approach, we preformed histology on lung samples of Lp-primed and unprimed mice 14 days post-Mtb infection and samples were scored by a blinded, trained pathologist. At this time point, there was limited signs of inflammation or tissue damage in either group (SFig 5, STable 2).

### Prior Mtb infection does not impact clearance of Lp

Mtb may impair effective innate immunity by suppressing innate immune responses [3, 4]. We sought to address this possibility by infecting mice with a ∼400 CFU dose of Mtb, allowing the infection to progress for 14 days, and then infecting these mice (or control uninfected mice) with Lp (Fig 4H). At 2 days post-Lp infection, there was no difference in Lp CFU between Mtb-infected and Mtb-uninfected hosts (Fig 4I). This experiment does not address local or cell-intrinsic immunosuppressive effects as Lp and Mtb are unlikely to infect the same cells but does suggest that Mtb does not induce a globally immunosuppressive environment in the lung.

### Single cell RNA-sequencing reveals changes to myeloid cell compartment after Lp-priming

Flow cytometry and cytokine analysis failed to reveal large, persistent changes to the lung after bacterial priming. As an alternative approach to characterize the effect of bacterial priming on innate immune responses to Mtb, we performed scRNA-seq on myeloid cells isolated from the lungs of unprimed and Lp-primed mice infected for 14 days with Mtb at an initial dose of ∼50 CFU (Fig 5A). CD64-positive cells were magnetically enriched, sort purified, and processed for library generation with the 10X Genomics platform. Mtb-infected and uninfected bystander cells were sorted. However, at this low initial dose and early time point, there were insufficient numbers of Mtb-infected cells to perform scRNA-seq. Therefore, we solely analyzed bystander cells. Protein expression was measured concurrently with mRNA by employing CITE-seq, allowing for weighted nearest neighbor (WNN) analysis to cluster cells on mRNA and protein expression and WNN uniform manifold approximation and projection (wnn.UMAP) reductions for data visualization (SFig 6A, B) [43]. Confirming the efficiency of our enrichment procedure, most cells isolated were myeloid cells.

**Fig 5.**
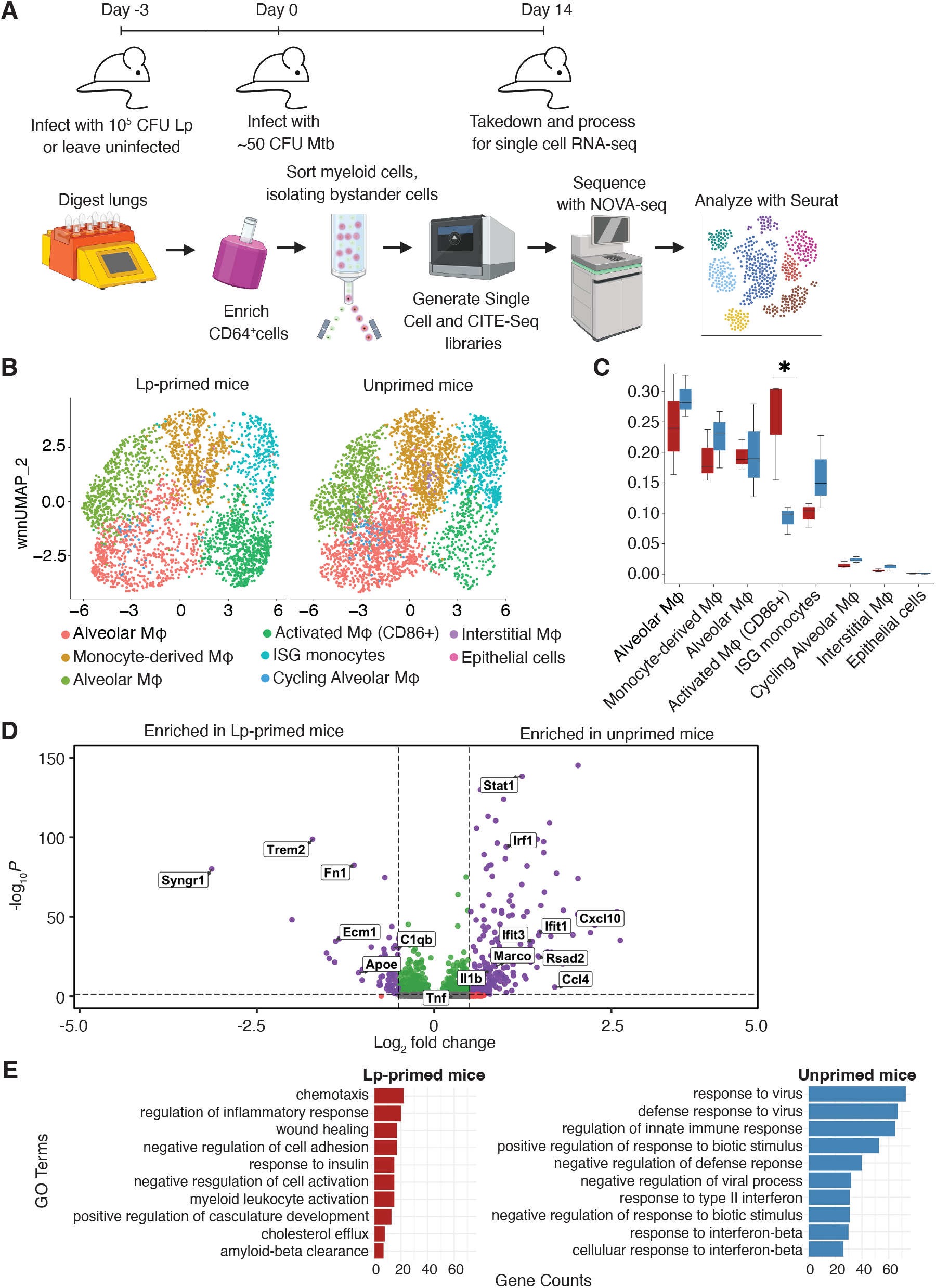
Lp priming alters gene expression and cell types in the lung after Mtb infection. (**A**) Model of the processing steps involved in generating the scRNA-seq dataset. (**B**) UMAP of unbiased clusters of myeloid cells in either unprimed or Lp-primed mice infected with Mtb. Only bystander cells are displayed. Lungs were analyzed 14 days post Mtb-mCherry infection. (**C**) Relative differences in cell proportions for each cluster between unprimed and Lp-primed samples. Significant differences were calculated with scCODA. (**D**) Differential gene expression (DEG) of bystander myeloid cells from Lp-primed or unprimed mice infected with Mtb. (**E**) DEGs were analyzed for gene functional annotations of GO terms.

Our scRNA-seq analysis indicated that unprimed and Lpprimed mice were generally similar. All clusters were represented in both datasets. Only one cluster had statistically significant shifts in cluster composition [44]. Lp-primed mouse lungs contained significantly higher numbers of activated macrophages, while unprimed mice exhibited elevated Interferon-stimulated gene (ISG)-expressing monocytes, although this change did not reach significance (Fig 5C) [45]. Examination of differential gene expression of all the cells in the dataset revealed that unprimed mice significantly upregulated genes associated with type I and type II interferon (IFN) signaling, including *Isg15, Cxcl10*, and IFIT family members (Fig 5D, SFig 6C, D). By contrast, Lp-primed mice induced genes associated with inflammation, but also control of inflammation such as *Apoe* and *Trem2* (Fig 5D, E, SFig 6D) [46].

Aligning with the above results, a Gene Ontology (GO) term analysis comparing GO terms differentially expressed between Lp-primed and unprimed mice, revealed distinct biological programs (Fig 5E). Cells from unprimed mice were enriched for GO terms associated with both type I and type II IFN, such as “response to virus” and “response to type II IFN.” By contrast, cell from Lp-primed mice were enriched for pathways associated with regulation of inflammation and cell migration, including “wound healing” and “chemotaxis.”

Overall, our data revealed that Lp infection results in persistent effects on the myeloid cell compartment that are evident even after 14 days of Mtb infection, well after Lp itself has been cleared. However, these differences in innate immune signaling do not dramatically alter the growth of Mtb bacteria, suggesting that Mtb replication is largely unaffected by a variety of anti-bacterial innate immune responses.

## Discussion

In mice, the initial *in vivo* innate immune response to Mtb fails to prevent the expansion of Mtb from a few bacilli to more than 10^5^ bacilli within 14 days (Fig 1) [24]. This rate of expansion corresponds to an approximate doubling time of 24h, similar to the doubling rate in rich media. In nonhuman primate infections, individual Mtb bacilli can also replicate largely unrestrained for several weeks [28, 47]. However, after the onset of adaptive immunity, innate immune clearance mechanisms—such as the microbicidal activity of macrophages—are critical for restriction of Mtb, as mice with genetic deficiencies in innate immunity eventually exhibit rapid decline as well as loss of bacterial control [9, 10, 36, 48]. Mtb control during the adaptive phase has largely been attributed to enhanced macrophage activation following the increase of IFN-γ production, which acts upon myeloid cells, by CD4^+^ T cells, although other pathways are also at play [49-52]. Patients with Mendelian Susceptibility to Mycobacterial Disease (MSMD) typically carry genetic mutations in IFN-γ-related pathways [53-55], further emphasizing the role of innate immunity in mediating successful adaptive immune responses to Mtb.

Human Mtb infections are also presumably poorly controlled by the innate immune system alone, though this is difficult to demonstrate experimentally. So-called “resisters”, who remain persistently negative by interferon-gamma release assay (IGRA) and tuberculin skin test (TST) despite exposure to Mtb, had been thought possibly to clear Mtb with innate immunity alone [56]. However, recent publications have demonstrated that these individuals likely control Mtb infections with alternative adaptive immune responses that were not detected by some diagnostic tests [57-59]. Studying the characteristics of the human innate immune response to Mtb poses challenges due to the difficulty in identifying recently infected individuals and assessing the localized responses within their lungs. Thus, genetically tractable mouse models provide an opportunity to examine the role of innate immune responses in control of Mtb replication or clearance.

In this study, we examined alternative Mtb infection models to see whether they might reveal innate immune mechanisms of Mtb control. We first employed a previously described ULD infection model [24]. ULD infections clearly demonstrated that the innate immune system alone is unable to clear or even control infections seeded by even 1-3 bacilli, indicating that the size of the initial infectious dose is not the limiting factor for innate immune control of Mtb *in vivo*. Wild-type mice and mice harboring mutations in key genes critical for Mtb control (e.g., *Ifngr1, Myd88, Atg5, Tnfr1*, and *Sp140*) had comparable CFU in the lungs at 14 days post ULD-infection, and the fraction of infected mice were not significantly different when comparing wild-type to immune-deficient animals. While we did not expect to see an effect due to the loss of IFN-γ signaling, as IFN-γ is thought to be mainly derived from CD4^+^ T cells in the context of Mtb infection, these other pathways are known to be critical to Mtb responses in innate immune cells [13, 36, 60-62].

A previous study of ULD infected mice immunized with BCG demonstrated that some ULD-infected mice have undetectable CFU burden at 14-15 days post-infection, suggesting that they control Mtb replication at early time points [29]. However, unvaccinated mice do not appear to harbor undetectable Mtb infection at this time point [29]. In contrast to BCG, which protects mice against Mtb disease, we do not expect mice lacking in key innate immune pathways to harbor undetectable Mtb bacteria. Further, while redundant mechanisms could mask early control of Mtb, the innate immune deficient mouse lines we selected cover a broad range of innate immune signaling pathways. These results do not rule out the possibility that innate immune cell activation, differentiation, survival or migration might be altered in the knockout mice; we can only conclude that even if such differences exist, they do not have a significant impact on the ability of the innate immune system to restrict bacterial burdens. Our results suggest that genes known to play important roles in Mtb control during the adaptive phase do not exert substantial effects during the innate phase of the infection, even at highly limited infectious doses.

We attempted to enhance innate immune responses to Mtb by pet shop mouse co-housing. Pet-shop co-housed animals have previously been shown to exhibit enhanced control of *Listeria* [31]. However, co-housed mice failed to control Mtb replication in the lung, suggesting that even the innate immune response of primed animals is insufficient to control Mtb. However, pet shop co-housing experiments are variable and technically complex. Thus, we turned to Lp co-infection as an alternative approach for priming innate immunity. We reasoned that since Lp elicits a highly effective anti-bacterial innate immune response, this response could inhibit Mtb replication. Use of Lp also negated the possibility of inducing Mtb-specific adaptive immune responses that might confound the results. However, Lp co-infections also failed to substantially enhance innate immune control of Mtb replication in the lung. Together, these data lead us to conclude that innate immunity exhibits a limited capacity to control Mtb replication.

Interestingly, we found the Lp priming reduced Mtb burdens in the spleen at 14 days post-infection. Dissemination of Mtb from the lung to extrapulmonary sites during the innate phase of the infection is not well understood. The mechanism by which Lp priming inhibits Mtb dissemination or replication in the spleen remains unclear from our data. However, we speculate that type I IFN signaling may play a role, as type I IFN can antagonize IFN-γ activity, and IFN-γ is critical to control of Mtb dissemination during the adaptive phase of infection [51, 63]. It is possible that elevated type I IFN in unprimed mice may suppress IFN-γ responses and permit greater dissemination in unprimed mice as compared to Lp-primed mice.

Mtb infection alone also induced inflammatory innate immune responses. It was possible that Lp priming did not greatly affect Mtb clearance because Lp priming did not enhance or change innate immune responses over and above those elicited by Mtb alone. Indeed, we demonstrated that by 14 days post-infection, the cellular innate immune response to Mtb in naïve and Lp-primed mice was generally similar, characterized by the presence of neutrophils, monocytes, and macrophages to the lung, suggesting that the cellular response by this time point may be driven largely by the innate immune response to Mtb. Since our analysis was limited to the 14 day post-infection time point, it did not capture the possible effects of Lp priming on the immune response in the hours and early days post-Mtb infection. Additionally, as our data set does not include Mtb-infected cells, we could not analyze cell-specific roles on control of Mtb replication at a single cell level. However, our scRNA-seq experiment revealed some notable differences in cell populations and interferon responses between Lp-primed and unprimed mice. While this scRNA-seq experiment captured only uninfected cells, we were able to uncover durable impacts of Lp priming.

Lp-primed mice exhibited an increase in activated macrophages expressing CD86, a cell-surface protein that provides co-stimulation to T cells [64, 65]. Interestingly, CD86 is reduced in TB patients compared to healthy controls [66], though given its role in T cell co-stimulation, it is likely that CD86 acts primarily via adaptive rather than innate immunity. In addition to CD86^+^ macrophages, Lp-primed mice expressed genes associated with resolution of inflammation and wound healing (Fig 5D, E). With the infection of mice with Lp for 3 days prior to 14 days of Mtb infection, Lp primed mice have been infected for a total of 17 days, versus only 14 days for the Mtb-only infected mice. The differences we see in Lp-primed mice may reflect a more advanced progression of the anti-bacterial immune response and a shift towards disease tolerance in the primed mice versus unprimed mice.

Our scRNA-seq also revealed that unprimed, Mtb-infected mice possessed a population of ISG-expressing macrophages and showed upregulated ISG expression compared to Lp-primed mice. Many ISGs are upregulated in response to both type I IFN and type II IFN, but we suspect that type I IFN signaling may drive this ISG signature based on the results from our GO analysis [60, 63]. In both mice and humans, type I IFN production can be associated with progression of TB, preceding active infection and predictive of poor clinical outcomes in humans [67, 68]. The mechanism of dampened ISG induction following Lp priming compared to unprimed, Mtb-infected mice, despite the known ability of Lp to induce ISG expression, remains unclear, but may reflect an immunoregulatory feedback loop [69, 70].

The differences we see in Lp-primed mice may reflect a more advanced progression of the anti-bacterial immune response and a shift towards disease tolerance in the primed mice versus unprimed mice. Disease tolerance plays a critical role as strategy to defend the host against the damaging impacts of infection [71, 72]. In wild-type C57BL/6 mice, Mtb infection is characterized by a balance of protective versus detrimental inflammation, but eventually results in progressive inflammatory damage to the lung tissue [73]. The transcriptional changes we observe by scRNA-seq could reflect alterations in disease tolerance even if they do not result in changes in Mtb burdens. We attempted to evaluate whether innate priming by Lp influenced tissue pathology, but at the day 14 timepoint there was very limited pathology and thus it was unclear whether differences in the innate response affected disease tolerance. Analysis at later timepoints would be confounded by potential effects of adaptive immunity.

Taken together, our data demonstrate the robust resistance of Mtb to innate immunity under diverse experimental conditions. Although Mtb may evade or impair the induction of specific innate immune pathways, the innate response to Mtb is overall robust and is characterized by the recruitment of cells and induction of anti-bacterial innate responses that are highly effective against other lung pathogens, such Lp. We therefore propose that a major reason for the failure of innate immunity to control Mtb is the robust resistance of Mtb to cellular stress and damage. Mtb exhibits resilience against antimicrobial effectors such as reactive oxygen, nitrogen species, and aldehydes, likely through intrinsic mechanisms such as its thick and waxy mycomembrane, a slow replication rate, and effective repair systems [74-76]. These characteristics have earned Mtb the epithet “the honey badger of pathogens” [77]. Like the honey badger that famously resists cobra venom or porcupine quills, Mtb appears impervious to the anti-bacterial arsenal that can rapidly clear other infectious agents [78]. This exceptional resistance to innate immune effectors may be an important factor contributing to the extraordinary success of Mtb as a human pathogen.

## Materials and Methods

### Ethics Statement

All procedures involving the use of mice were approved by the University of California, Berkeley Institutional Animal Care and Use Committee (Protocol Number 2014-09-6665-2). All protocols conform to federal regulations, the National Research Council’s Guide for the Care and Use of Laboratory Animals and the Public Health Service’s Policy on Humane Care and Use of Laboratory Animals.

### Animals

Mice were maintained under specific pathogen free conditions and housed at 23°C with a 12-hour light-dark cycle in accordance with the regulatory standards of the University of California Berkeley Institutional Animal Care and Use Committee. All mice were 6-12 weeks old at the start of procedures. C57BL/6, B6.129S7-*Ifngr1*^tm1Agt^/J *(Ifngr1*^*–/–*^), and B6.129S-*Tnfrsf1b*^tm1Imx^ *Tnfrsf1a*^tm1Imx^/J *(Tnfr*^*–/–*^) mice were purchased from Jackson Laboratories. *Atg5*^*fl/fl*^*-LysM*^*Cre*^ and *Atg5*^*fl/ fl*^ and mice were provided by Jeffery Cox at the University of California, Berkeley. *Myd88*^*–/–*^*Trif*^*–/–*^ mice were provided by Gregory Barton at the University of California, Berkeley. *Sp140*^*–/–*^ mice were previously generated by the Vance Lab [35].

### Pet shop co-housed mice

6-week-old female C57BL/6 mice were purchased from Jackson Laboratories and housed in Innovivo IVC Rat Cages. 8 C57BL/6 mice were housed per cage. Each group was co-housed with one female pet shop mouse purchased from the East Bay Vivarium (Berkeley, CA) for 60 days. Upon importation of the pet shop mice, each mouse was tested for serological presence of pathogens by Vrl Animal Health Diagnostics. Mice were weighed every 3-7 days and monitored for weight loss. When sufficient numbers of mice were available, we distributed mice from the same cage across experimental groups to allow comparison of cage mates to different pathogens.

### Bacterial strains

Mtb strain Erdman was a gift from Sarah Stanley at University of California, Berkeley. Mtb expressing Wasabi or mCherry were generated as previously described [60]. *Legionella pneumophila* strains were JR32 and isogenic mutants that were produced as previously described [41]. Lp expressing mCherry was produced and cultured as previously described [23]. The mCherry expressing plasmid was a gift from Sunny Shin [79]. All infections were performed with strains lacking *flaA*.

### *Mycobacterium tuberculosis* infections

For standard low-dose infections, a previously frozen aliquot of Mtb was diluted in 20 mL of distilled water. 9 mL of the diluted culture was loaded into the nebulizer of an inhalation exposure system (Glas-Col, Terre-Haute, IN) to deliver ∼10-50 CFU of bacteria per mouse as determined by measuring CFU in the lungs 1 day post-infection. For ultra-low dose infections, the infectious dose was first estimated by calculating the volume needed to deliver 1 CFU based on the volume needed to achieve an infectious dose of 20 CFUs in previous experiments. The frozen stock was serially diluted, and the appropriate volume was added to 20 mL of distilled water. The ultra-low infectious dose was confirmed by calculating the percent of infected mice (at 14 days post-infection) per experiment instead of measuring CFU in the lungs at 1 day post-infection [38]. For intranasal infections, a frozen stock of Mtb was grown in 7H9 buffer containing 0.05% Tween-80. At log phase, bacteria were subject to low-speed centrifugation and gentle sonication to achieve a single cell suspension. An average of 40 CFU were delivered to each mouse after anesthetization with ketamine and xylazine by intraperitoneal injection. To enumerate CFU at 1 day post-infection, lungs were homogenized in 1 mL of PBS containing 0.05% Tween-80 (PBS-T) and the entirety of the lung was plated across 4 plates. For other time points, 100 μL were taken from suspensions from lungs either collected in 1 mL of PBS-T or from the 5 mL RMPI solution used for flow cytometry sample collection. Samples were then serially diluted in PBS containing 0.05% Tween-80. Serial dilutions were plated on 7H11 plates supplemented with 0.5% glycerol and BBL Middlebrook OADC Enrichment media (BD Biosciences). Colonies were counted after 3 weeks incubation at 37°C.

### *Legionella pneumophila* infections

For infections, frozen cultures were streaked out onto BCYE plates and incubated at 37°C to obtain single colonies, which were subsequently streaked onto a new BCYE plate and incubated at 37°C to obtain a 1 cm by 1 cm bacterial patch. The patch was then collected and grown for ∼20 hours in CYE media with streptomycin (100 μg/mL). The optical density was measured at 600 nm. The culture was diluted to 2.5 × 10^6^ CFU/mL in sterile PBS. The mice were anesthetized with ketamine and xylazine by intraperitoneal injection the infected intranasally with 40 μL of inoculum to deliver 1 × 10^5^ CFU per mouse. The infectious dose was confirmed by measuring CFU in the lungs 4 hours post-infection. Lungs were harvested in either 5 mL of sterile water at the 4 hour time point or 1 mL of sterile water at all other time points. Samples were then serially diluted in water and plated on BCYE plates.

### *Listeria monocytogenes* infections

Lm infections were performed as previously described [80]. Briefly, mice were infected with strain 10403S via tail vein injection. 5 × 10^4^ CFU of Lm were delivered per mouse. The spleen and liver was harvest from each infected mouse at 48 hours post-infection.

### Cytokine Analysis

We used the LEGENDPlex Mouse Inflammation Panel (13-Plex) (BioLegend) to analyze expression of inflammatory cytokines in the lung. Mice were either primed with Lp or left unprimed and then infected with Mtb as described above. At 14 or 28 days post-infection, the lungs were harvested in 1 mL of PBS containing protease inhibitors (Roche), homogenized, and frozen. Samples were then thawed and removed from the BSL3 after decontamination by 2 rounds of centrifugation through a 0.2 μM filter microcentrifuge tubes. Samples were stained for cytokine expression following the manufacture’s kit and analyzed on an Aurora (Cytek) cytometer.

### Flow cytometry

For flow cytometry, mouse lungs were harvested at 14 days post-Mtb-infection. The entire lung was harvested into a gentleMACS C tube (Miltenyi Biotec) containing 3 mL of RPMI media with 70 μg/mL of Liberase TM (Roche) and 30 μg/mL of Dnase I (Roche). Lungs were processed with the lung_01 setting on the gentleMACS (Miltenyi Biotec) and incubated at 37°C for 30 minutes. To obtain a single cell suspension, the samples were then homogenized on lung_02 setting on the gentleMACS. 2 mL of phosphate-buffered saline (PBS) containing 20% Newborn Calf Serum (Thermo Fisher Scientific) was then added to quench the digestion. Samples were then filtered through 70 μM SmartStrainers (Miltenyi Biotec). The single cell suspension was then spun down and resuspended in 500 μL of PBS containing 5% Newborn Calf Serum and 0.05% sodium azide. 100 uL were then stained with antibodies for 30 minutes at room temperature with the following antibodies: BV421-labeled PD-L1 (HIH5, BD Biosciences), BV480-labeled B220 (RA3-6B2, BD Biosciences), CD90.2-labeled (53-2.1, BD Biosciences), SB645-labeled MHC Class II (I-A/I-E) (M5/114.15.2, Thermo Fisher Scientific), BV786-labeled Ly-6C (HK1.4, BioLegend), PE-labeled MerTK (DS5MMER, Thermo Fisher Scientific), APC-labeled CD64 (X54-5/7.1, BioLegend), APC-R700-labeled Siglec-F (E50-2440, BD Biosciences), APC/Fire 750-labeled Ly-6G (1A8, BioLegend), BUV396-labeled CD11b (M1/70, BD Biosciences), BUV496-labeled CD45 (30-F11, BD Biosciences), BUV737-labeled CD11c (HL3, BD Biosciences), and BV-785-labeld CD86 (GL-1, BioLegend). All staining cocktails contained True-Stain monocyte blocker (BioLegend), Super Bright Complete Staining Buffer (Thermo Fisher Scientific), and GhostDye 510 (Tonbo). Samples were then fixed for 30 minutes with Cytofix/Cytoperm (BD Biosciences) before removal from the BSL3. Cell numbers were calculated by adding CountBright Plus Absolute Counting Beads (Invitrogen) to each sample. Samples were analyzed on a LSRFortessa X-20 (BD Biosciences) or an Aurora (Cytek) cytometer. Data were analyzed using Flowjo Version 10 (BD Biosciences). Gating was performed as previously described [60]. Sorting Immune Cells for single cell (sc)RNA-seq 3 C57BL/6 mice were infected with 10^5^ CFU of Lp-Δ*flaA* expressing an mCherry plasmid. After 3 days, these mice, along with 3 other naïve C57BL/6 mice, were subsequently infected with ∼50 CFU of Mtb-mCherry. At 14 days post-infection, the lungs were harvested and processed as for flow cytometry, as described above. Samples were then stained on ice for 30 minutes with APC-labeled CD64 (X54-5/7.1, BioLegend). Myeloid cells were magnetically enriched using an EasySep APC Positive Selection Kit II (StemCell Technologies) and MojoSort Magnets (BioLegend). Samples were then stained on ice for 45 minutes with the following anti-mouse antibodies in the presence of True-Stain Monocyte Blocker (BioLegend): TruStain FcX PLUS (S17011E, BioLegend), BV786-labeled CD45.2 (104, BioLegend), and Pacific Blue-labeled B220 (RA3-6B2, BioLegend), CD90.2 (53-2.1, BioLegend), and Ly6G (1A8, BioLegend). The Pacific Blue-labeled antibodies were used to exclude neutrophils, T cells, and B cells. Additionally, the following TotalSeq-A-labeled anti-mouse antibodies were added to detect protein expression in the scRNA-seq dataset: Ly6C (HK1.4, BioLegend), CD274 (MIH6, BioLegend), Siglec F (S17007L, BioLegend), CSF1R (AFS98, BioLegend), CD11b (M1/70, BioLegend), CD86 (GL-1, BioLegend), and MHC II (M5/114.15.2, BioLegend). A unique anti-mouse TotalSeq-A Hashtag antibody (1-6; BioLegend) was added to each sample to allow up to 6 samples to be mixed per lane on the 10X Chromium Next GEM Chip (10X Genomics). After staining, cells were resuspended in PBS with 5% Newborn Calf Serum and Sytox Blue Dead Cell Stain (Thermo Fisher Scientific). Cells were sorted using a 120 μm microfluidic sorting chip in a 4 laser SH-800 cell sorter (Sony) on the purity setting. Cells were collected in PBS containing 50% fetal calf serum.

### scRNA-seq library generation and sequencing

Single-cell RNA-sequencing was preformed following previously published protocols [60]. Briefly, the v3.1 chemistry Chromium Single Cell 3’ Reagent Kit (10X Genomics) was used to generate scRNA-seq libraries. The manufacturer protocol was followed, with the following minor modifications. Lanes of the 10X Chromium Next GEM Chip were super-loaded with 29,000 cells. During the loading step, 0.5 U/μL RNaseOUT Recombinant Ribonuclease Inhibitor (Invitrogen) was added to single cell RT master mix. During the cDNA amplification, 1 μL of ADT and HTO additive primers (0.2 μM stock) were added as recommended by the CITE-seq and Cell Hashing Protocol [43]. After the purification step, samples were decontaminated by 2 rounds of centrifugation through a 0.2 μM filter microcentrifuge tubes and removed from the BSL3. cDNA libraries were prepared following the 10X Genomics protocol. The ADT and HTO libraries were prepared following the CITE-Seq and Cell Hashing Protocols. DNA quality was measured using a TapeStation (Agilent). The cDNA, ADT and HTO libraries were pooled at the following proportion: 80% cDNA, 10% ADT, and 10% HTO. The libraries were sequenced on an Illumina platform (Azenta).

### Single cell RNA-seq data processing

Raw reads were processed into count matrices with CellRanger v8.0.0 (10x Genomics). The matrices were analyzed in R (RStudio) using Seurat v5.1.0. Cells were filtered to retain singlets expressing 200–4,500 genes with <5% mitochondrial reads. RNA data were normalized with NormalizeData and subsequently transformed with SCTransform while regressing out mitochondrial content. Principal component analysis (PCA) and UMAP were performed using the top 30 dimensions, and nearest-neighbor graphs were built on the same dimensions. ADT features were set as variable, normalized using centered log-ratio transformation with a margin of 2, scaled, and reduced by PCA using three principal components. Multi-modal integration was performed using FindMultiModalNeighbors, combining five RNA dimensions and three ADT dimensions. UMAP was then computed using weighted nearest-neighbor graphs, and clustering was performed with FindClusters (algorithm = 3; resolution = 0.3). Data visualization was performed with tidyverse, EnhancedVolcano, and Seurat plotting functions. scCODA was used to analyze cluster composition using the alveolar macrophage cluster (Cluster 0) as a reference population [44]. Gene Ontology analysis was used to identify biological processes enriched in each experimental group by comparing differentially expressed genes between unprimed and Lp-primed mice [81, 82]. Default parameters in Seurat were used. Genes were submitted to Gene Ontology Biological Process enrichment analysis using the clusterProfiler package, with the mouse genome annotation database as a reference [83]. Redundant terms were removed using the simplify() function (cutoff = 0.7).

## Data Accessibility

Single cell RNA-sequencing data is available from the NCBI Gene Expression Omnibus repository under accession number GSE328967. The code for analysis is available on Github: https://github.com/marianrfairgrieve/HoneyBadger.

## Author Contributions

Conceptualization: MRF, DIK, REV; Data curation: MRF; Formal analysis: MRF; Investigation: MRF, ECB, RAC, DIK; Methodology: MRF; Validation: MRF; Visualization: MRF; Writing–original draft: MRF; Writing–review and editing: REV; Funding Acquisition, Resources, Supervision: REV.

## Acknowledgments

We thank members of the Vance, Barton, Stanley, and Cox labs for advice and discussion and Victoria Chevée and Andrea Anya-Sanchez for assistance with Listeria infections. We also thank Jesse Rodriguez, Joceline Morales, the East Bay Vivarium, and the Office of Laboratory Animal Care for assistance with mice. MRF was supported by a Sidney Mac Donald Russell Family fellowship from the UC Berkeley Center for Emerging and Neglected Diseases. Research in the laboratory of REV is funded by an Investigator Award from the Howard Hughes Medical Institute and by NIH grants AI066302, AI075039, and AI155634. The funders had no role in study design, data collection and analysis, decision to publish, or preparation of the manuscript. Model figures were created in BioRender.

---

**Supplementary Figure 1.**
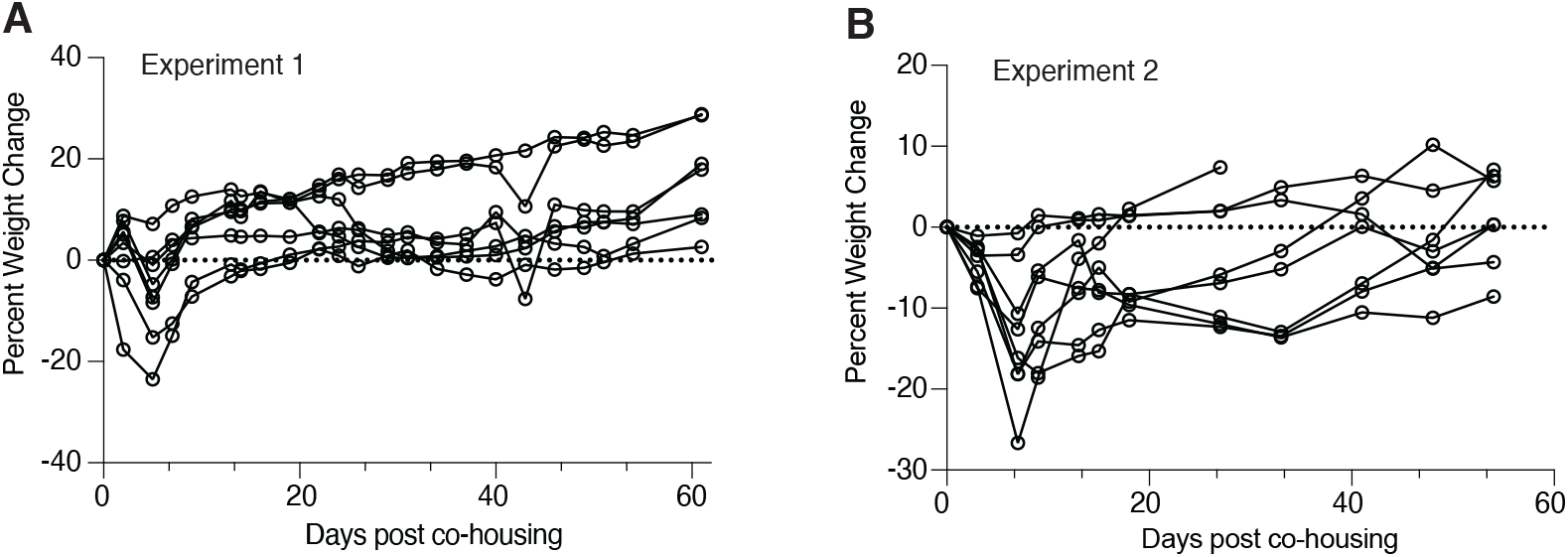
Weight changes in co-housed mice. (**A-B**) Average percent weight change of C57BL/6 mice cohoused with pet shop mice in two individual experiments. Each line represents the average weight change of up to 8 mice in one cage. Mice were weighed every 3-7 days over 60 days.

**Supplementary Fig 2.**
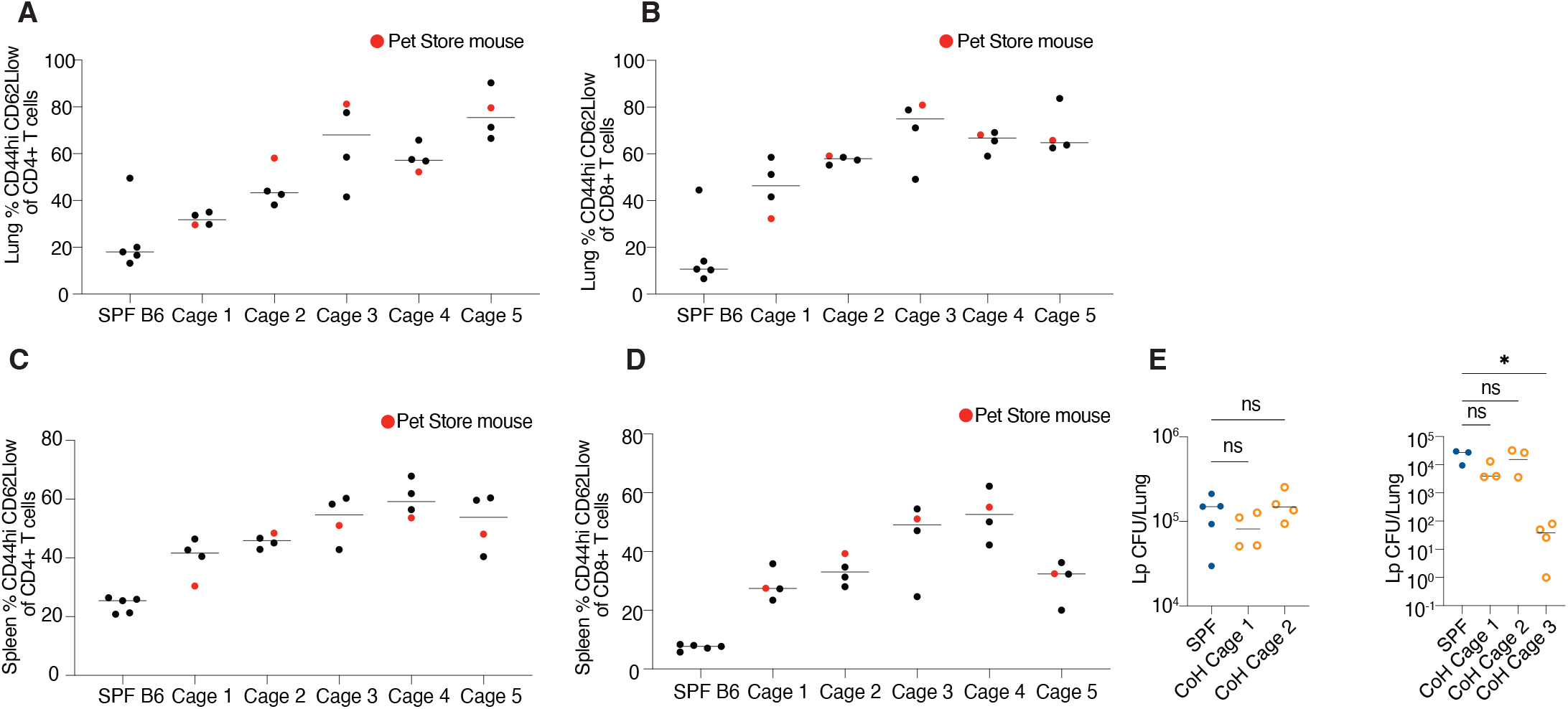
The founder pet shop mouse influences cell counts and infection outcomes in cohoused C57BL/6 mice. (**A-B**) Percent of CD4^+^ (A) or CD8^+^ (B) T effector memory cells in the lungs of either SPF C57BL/6 mice, pet shop cohoused, or pet shop mice (red). (**C-D**) % of CD4^+^ (C) or CD8^+^ (D) T effector memory cells in the spleens of either SPF C57BL/6 mice, pet shop co-housed, or pet shop mice (red). (**E**) Lp lung CFU from two separate experiments at 2 days post infection. The bars represent the median where CFU was detected. Mann-Whitney tests were used to calculate significance. *p ≤ 0.05.

**Supplementary Fig 3.**
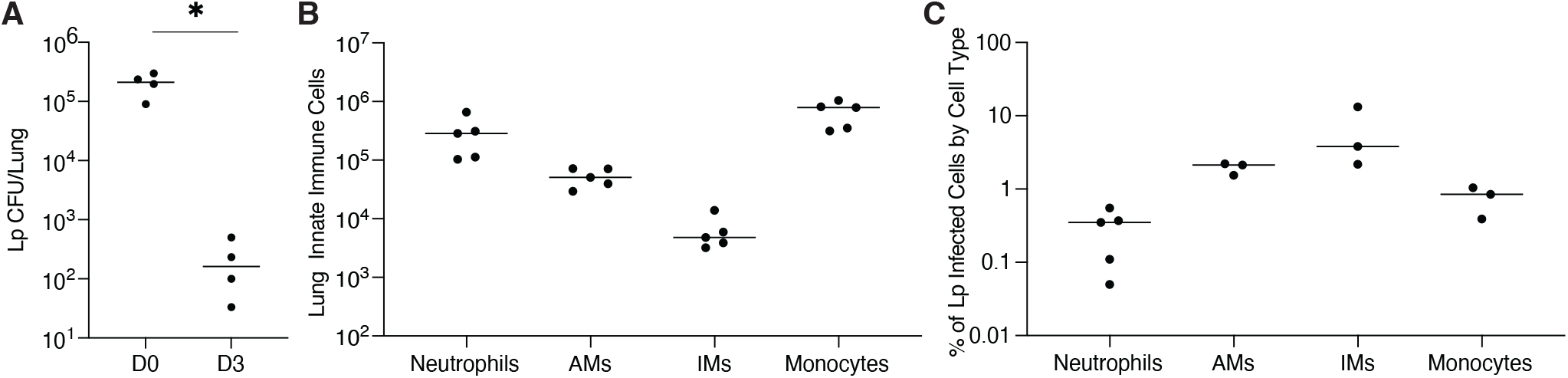
Characterizing Lp infection in C57BL/6 mice. (**A**) Lp Lung CFU measured 4 hours post infection (D0) and 3 days post-infection (D3) (**B**) Lung innate immune cells 1 day post-Lp infection. (C) Localization of Lp as measured by the detection of fluorescent bacteria as a percent of each cell type at 2 days post-infection.

**Supplementary Fig 4.**
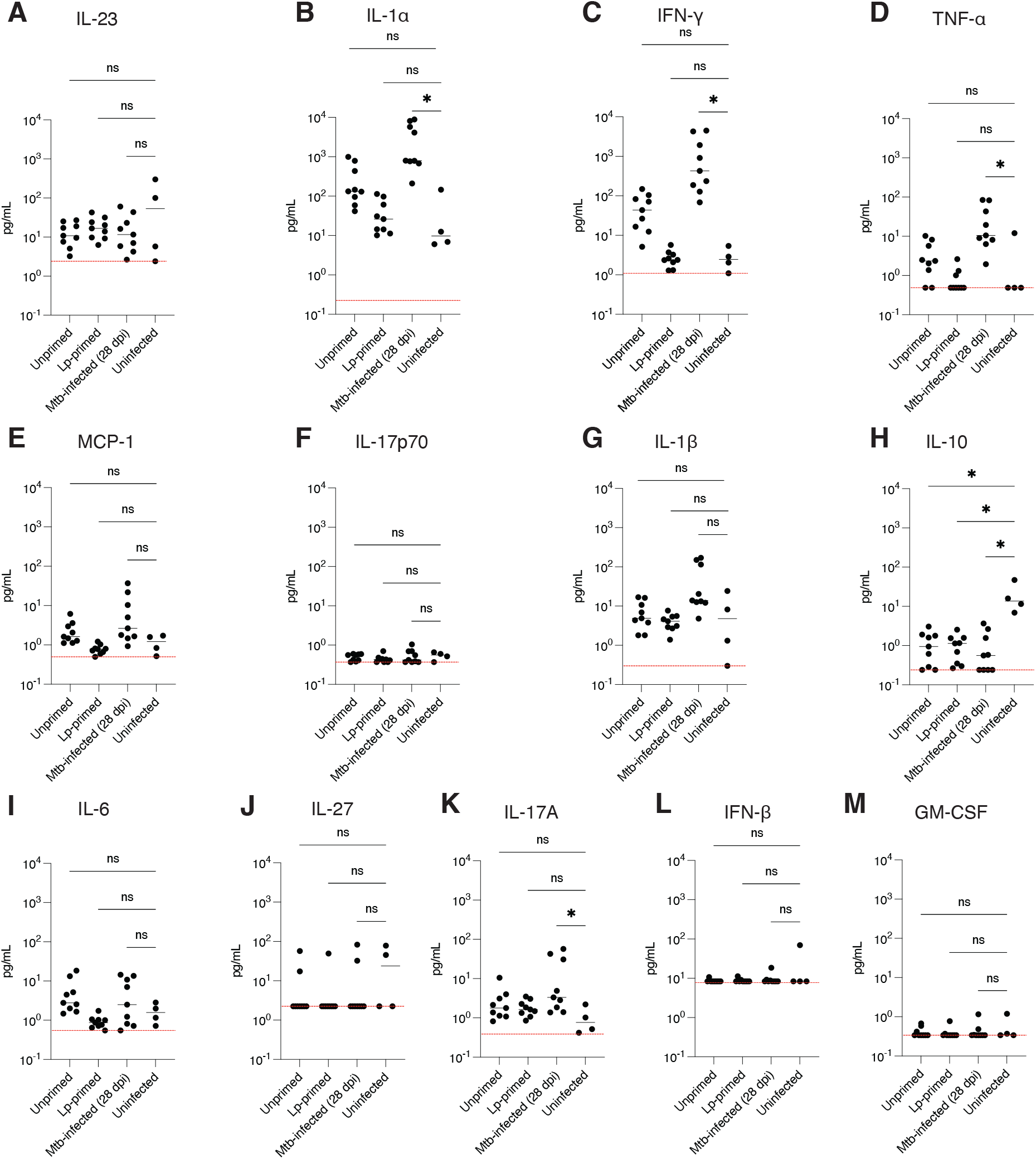
Cytokine expression in the lungs of unprimed and Lp-primed C57BL/6 mice. (**A-M**) Quantification of cytokine expression measured by LEGENDplex in unprimed or Lp-primed mice after 14 days of Mtb infection. As a positive control, mice infected with Mtb for 28 days were included. (A) IL-23, (B) IL-1α, (C) IFN-γ, (D) TNF-α, (E) MCP-1, (F) IL-17p70, (G) IL-1β, (H) IL-10, (I) IL-6, (J) IL-27, (K) IL-17A, (L) IFN-β, (M) GM-CSF. Results shown are from 2 combined experiments. Significance was calculated with a Kruskal-Wallis test followed by Dunn’s multiple comparisons test. *p ≤ 0.05.

**Supplementary Fig 5.**
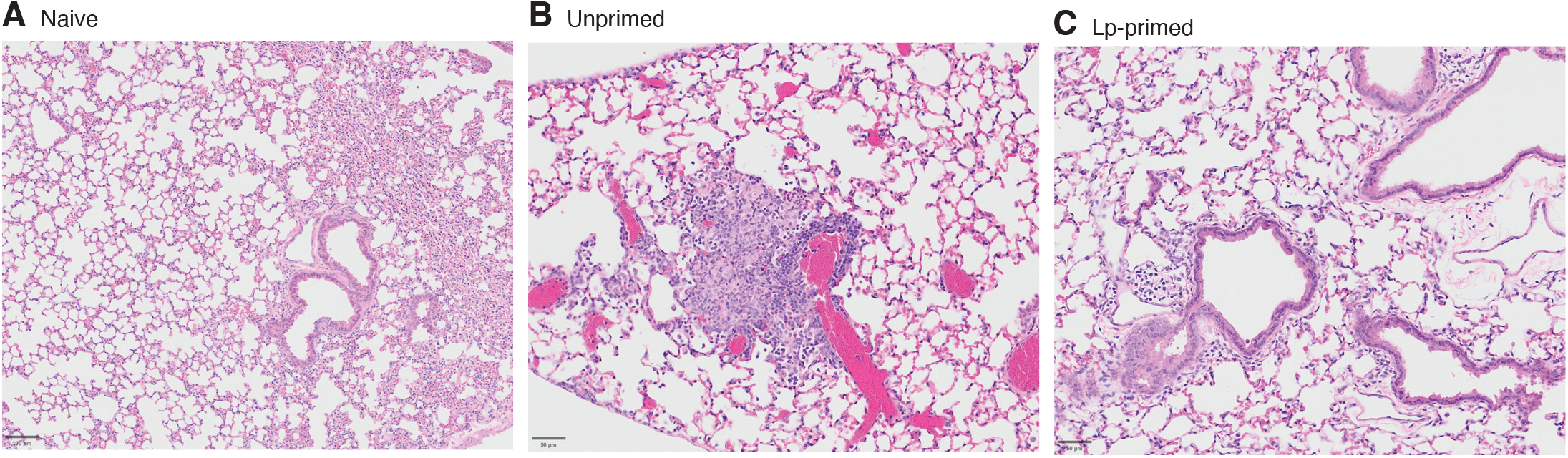
Histology and acid-fast staining of lungs. (**A-C**) Representative H&E images of either naïve, Lp-primed, or unprimed mice.

**Supplementary Table 1.** Pathogen diagnostic panel of pet shop mice. Pathogen diagnostic panel performed on blood samples of pet shop mice by serology or immunofluorescence assay (IFA) over two independent experiments. 7 mice were tested for each experiment.

| AGENTS TESTED | Methodology | Experiment 1 |  |  | Experiment 2 |  |  |
| --- | --- | --- | --- | --- | --- | --- | --- |
|  |  | Tested | Positive | Negative | Tested | Positive | Negative |
| Mouse Parvovirus MPV | SEROLOGY | 7 | 3 | 4 | 7 | 2 | 5 |
| Mouse Minute virus | SEROLOGY | 7 | 0 | 7 | 7 | 4 | 3 |
| Parvo NS-1 | SEROLOGY | 7 | 4 | 3 | 7 | 3 | 4 |
| Mouse Hepatitis MHV | SEROLOGY | 7 | 6 | 1 | 7 | 7 | 0 |
| Murine Norovirus MNV | SEROLOGY | 7 | 2 | 5 | 7 | 3 | 4 |
| Theilovirus(GD7TMEV) | SEROLOGY | 7 | 0 | 7 | 7 | 0 | 7 |
| EDIM/Rotavirus | SEROLOGY | 7 | 0 | 7 | 7 | 3 | 4 |
| Sendai virus | SEROLOGY | 7 | 0 | 7 | 7 | 0 | 7 |
| Pneumonia virus PVM | SEROLOGY | 7 | 0 | 7 | 7 | 2 | 5 |
| Reovirus 1,2,3 | SEROLOGY | 7 | 0 | 7 | 7 | 0 | 7 |
| LCMV | SEROLOGY | 7 | 0 | 7 | 7 | 0 | 7 |
| MAD-1 (MAV-FL) | SEROLOGY | 7 | 0 | 7 | 7 | 0 | 7 |
| MAD-2 (K87) | SEROLOGY | 7 | 0 | 7 | 7 | 0 | 7 |
| Ectromelia virus | SEROLOGY | 7 | 0 | 7 | 7 | 0 | 7 |
| Polyoma virus | SEROLOGY | 7 | 0 | 7 | 7 | 0 | 7 |
| Murine CMV | IFA | 7 | 0 | 7 | 7 | 0 | 7 |
| Carbacillus CARB | SEROLOGY | 7 | 0 | 7 | 7 | 0 | 7 |

**Supplementary Table 2.** Histological analysis of naïve, unprimed, or Lp-primed mouse lungs. Sections of mouse lungs were stained and analyzed by a trained, blinded pathologist for signs of inflammation after 14 days of Mtb <u>infection.</u>

| Sample | Inflammatory cell aggregation, interalveolar and septal | Inflammatory cell infiltration, perivascular and peribronchial | Vasculitis, lymphohistiocytic |
| --- | --- | --- | --- |
| Naïve | - | - | - |
| Unprimed sample 1 | Mild, multifocal, multilobe | Mild, multifocal, multilobe | Mild, multifocal |
| Unprimed Sample 2 | Mild, multifocal, multilobe | Mild, multifocal, multilobe | Suspected, multifocal |
| Unprimed Sample 3 | Mild, multifocal, multilobe | Mild, multifocal, multilobe | Mild, multifocal |
| Lp-primed Sample 1 | - | Mild, multifocal, multilobe | - |
| Lp-primed Sample 2 | Minimal (extension from perivascular infiltrate) | Mild, multifocal, multilobe | Suspected, multifocal |
| Lp-primed Sample 3 | Minimal, focal | Minimal, multifocal, multilobe | - |

**Supplementary Fig 6.**
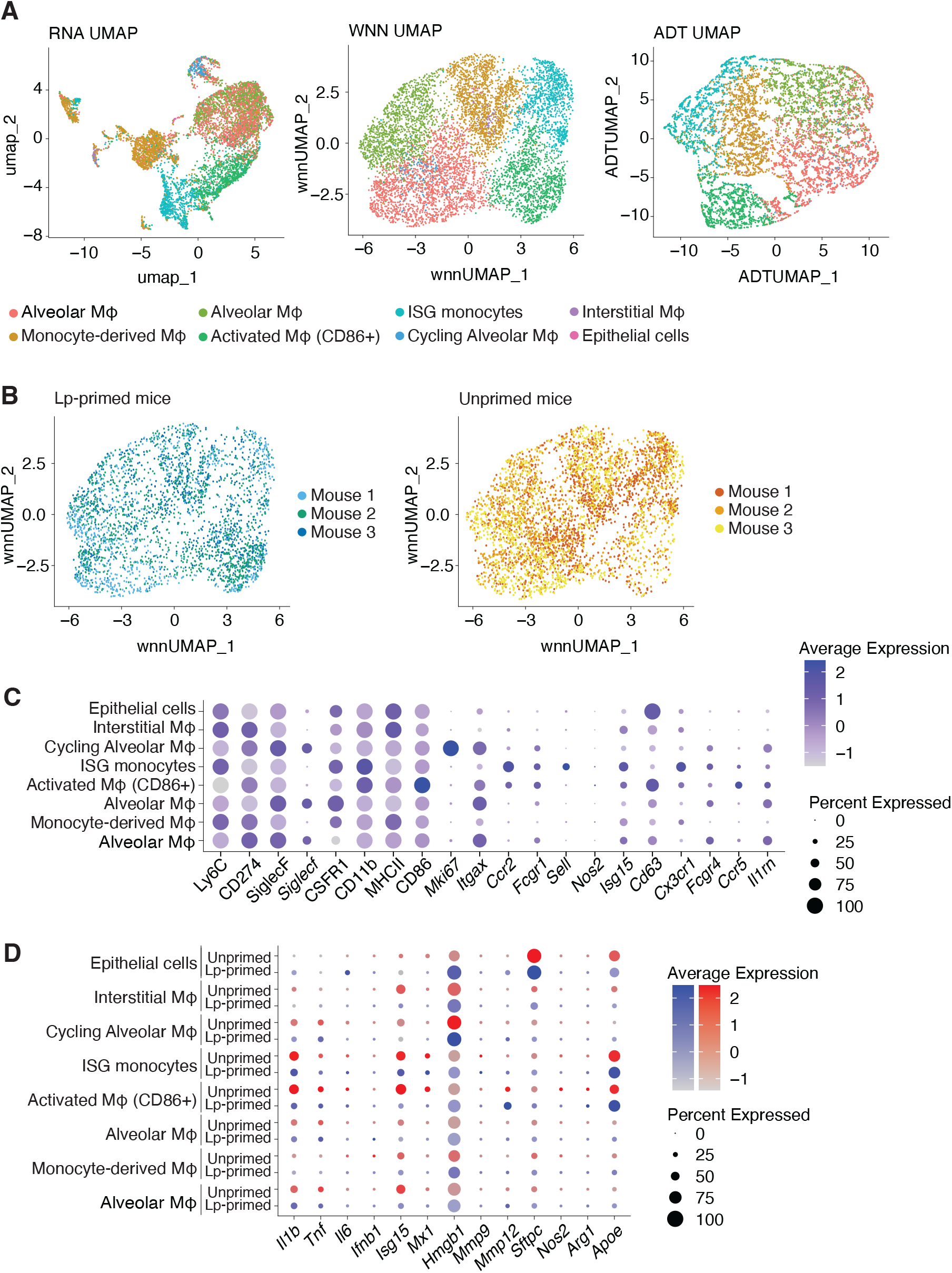
Identification of lung myeloid cells and markers of activation from scRNA-seq. **(A)** UMAP plots showing cell clustering of combined mRNA and protein expression, mRNA expression, or protein expression. Only bystander cells are shown. (**B**) UMAP plots showing cell clustering of biological replicates of unprimed or Lp-primed mice. Only bystander cells are shown. (**C**) Protein and mRNA expression of lineage-defining myeloid cell markers used for cluster classification. Both unprimed and Lp-primed mice were analyzed to identify clusters. (**D**) mRNA expression of cell state markers in unprimed and Lpprimed mice. Inflammatory mediators: *Il1b, Tnf, Il6*; Type I IFN and ISGs: *Ifnb1, Isg15, Mx1*; Macrophage activation: *Nos2, Arg1, Apoe*; Lung injury or dysfunction: *Hmgb1, Mmp9, Mmp12, Sftpc*.

## References

1. Ravesloot-Chávez MM, Van Dis E, Stanley SA. The Innate Immune Response to *Mycobacterium tuberculosis*. Annu Rev Immunol. 2021;39:611–37. Epub 20210226. doi: 10.1146/annurev-immunol-093019-010426. PubMed PMID: 33637017.

2. Cohen SB, Gern BH, Delahaye JL, Adams KN, Plumlee CR, Winkler JK, et al. Alveolar Macrophages Provide an Early *Mycobacterium tuberculosis* Niche and Initiate Dissemination. Cell Host Microbe. 2018;24(3):439–46.e4. Epub 20180823. doi: 10.1016/j.chom.2018.08.001. PubMed PMID: 30146391; PubMed Central PMCID: PMC6152889.

3. Rothchild AC, Olson GS, Nemeth J, Amon LM, Mai D, Gold ES, et al. Alveolar macrophages generate a noncanonical NRF2-driven transcriptional response to *Mycobacterium tuberculosis* in vivo. Sci Immunol. 2019;4(37). doi: 10.1126/sciimmunol.aaw6693. PubMed PMID: 31350281; PubMed Central PMCID: PMC6910245.

4. Pisu D, Johnston L, Mattila JT, Russell DG. The frequency of CD38+ alveolar macrophages correlates with early control of *M. tuberculosi*s in the murine lung. Nat Commun. 2024;15(1):8522. Epub 20241002. doi: 10.1038/s41467-024-52846-w. PubMed PMID: 39358361; PubMed Central PMCID: PMC11447019.

5. Pisu D, Huang L, Narang V, Theriault M, Lê-Bury G, Lee B, et al. Single cell analysis of *M. tuberculosis* phenotype and macrophage lineages in the infected lung. J Exp Med. 2021;218(9). Epub 20210722. doi: 10.1084/jem.20210615. PubMed PMID: 34292313; PubMed Central PMCID: PMC8302446.

6. Mai D, Jahn A, Murray T, Morikubo M, Lim PN, Cervantes MM, et al. Exposure to *Mycobacterium* remodels alveolar macrophages and the early innate response to *Mycobacterium tuberculosis* infection. PLoS Pathog. 2024;20(1):e1011871. Epub 20240118. doi: 10.1371/journal.ppat.1011871. PubMed PMID: 38236787; PubMed Central PMCID: PMC10796046.

7. Mata E, Tarancon R, Guerrero C, Moreo E, Moreau F, Uranga S, et al. Pulmonary BCG induces lung-resident macrophage activation and confers long-term protection against tuberculosis. Sci Immunol. 2021;6(63):eabc2934. Epub 20210924. doi: 10.1126/sciimmunol.abc2934. PubMed PMID: 34559551.

8. Wolf AJ, Desvignes L, Linas B, Banaiee N, Tamura T, Takatsu K, et al. Initiation of the adaptive immune response to *Mycobacterium tuberculosis* depends on antigen production in the local lymph node, not the lungs. The Journal of experimental medicine. 2008;205:105–15.

9. Scanga CA, Bafica A, Feng CG, Cheever AW, Hieny S, Sher A. MyD88-deficient mice display a profound loss in resistance to *Mycobacterium tuberculosis* associated with partially impaired Th1 cytokine and nitric oxide synthase 2 expression. Infect Immun. 2004;72(4):2400–4. doi: 10.1128/IAI.72.4.2400-2404.2004. PubMed PMID: 15039368; PubMed Central PMCID: PMC375220.

10. Bean AG, Roach DR, Briscoe H, France MP, Korner H, Sedgwick JD, et al. Structural deficiencies in granuloma formation in TNF gene-targeted mice underlie the heightened susceptibility to aerosol *Mycobacterium tuberculosis* infection, which is not compensated for by lymphotoxin. J Immunol. 1999;162(6):3504–11. PubMed PMID: 10092807.

11. Sugawara I, Yamada H, Li C, Mizuno S, Takeuchi O, Akira S. Mycobacterial infection in TLR2 and TLR6 knockout mice. Microbiol Immunol. 2003;47(5):327–36. doi: 10.1111/j.1348-0421.2003.tb03404.x. PubMed PMID: 12825894.

12. Bafica A, Scanga CA, Feng CG, Leifer C, Cheever A, Sher A. TLR9 regulates Th1 responses and cooperates with TLR2 in mediating optimal resistance to *Mycobacterium tuberculosis*. J Exp Med. 2005;202(12):1715–24. doi: 10.1084/jem.20051782. PubMed PMID: 16365150; PubMed Central PMCID: PMC2212963.

13. Di Paolo NC, Shafiani S, Day T, Papayannopoulou T, Russell DW, Iwakura Y, et al. Interdependence between Interleukin-1 and Tumor Necrosis Factor Regulates TNF-Dependent Control of *Mycobacterium tuberculosis* Infection. Immunity. 2015;43(6):1125–36. doi: 10.1016/j.immuni.2015.11.016. PubMed PMID: 26682985; PubMed Central PMCID: PMC4685953.

14. Kimmey JM, Huynh JP, Weiss LA, Park S, Kambal A, Debnath J, et al. Unique role for ATG5 in neutrophil-mediated immunopathology during *M. tuberculosis infection*. Nature. 2015;528(7583):565–9. Epub 20151209. doi: 10.1038/nature16451. PubMed PMID: 26649827; PubMed Central PMCID: PMC4842313.

15. Nair S, Huynh JP, Lampropoulou V, Loginicheva E, Esaulova E, Gounder AP, et al. Irg1 expression in myeloid cells prevents immunopathology during *M. tuberculosis* infection. J Exp Med. 2018;215(4):1035–45. Epub 20180306. doi: 10.1084/jem.20180118. PubMed PMID: 29511063; PubMed Central PMCID: PMC5881474.

16. Dorhoi A, Desel C, Yeremeev V, Pradl L, Brinkmann V, Mollenkopf HJ, et al. The adaptor molecule CARD9 is essential for tuberculosis control. J Exp Med. 2010;207(4):777–92. Epub 20100329. doi: 10.1084/jem.20090067. PubMed PMID: 20351059; PubMed Central PMCID: PMC2856020.

17. MacMicking JD, North RJ, LaCourse R, Mudgett JS, Shah SK, Nathan CF. Identification of nitric oxide synthase as a protective locus against tuberculosis. Proc Natl Acad Sci U S A. 1997;94(10):5243–8. doi: 10.1073/pnas.94.10.5243. PubMed PMID: 9144222; PubMed Central PMCID: PMC24663.

18. Peignier A, Kim J, Lemenze A, Parker D. Monocyte-regulated interleukin 12 production drives clearance of *Staphylococcus aureus*. PLoS Pathog. 2024;20(10):e1012648. Epub 20241017. doi: 10.1371/journal.ppat.1012648. PubMed PMID: 39418302; PubMed Central PMCID: PMC11521269.

19. Tsai WC, Strieter RM, Mehrad B, Newstead MW, Zeng X, Standiford TJ. CXC chemokine receptor CXCR2 is essential for protective innate host response in murine *Pseudomonas aeruginosa* pneumonia. Infect Immun. 2000;68(7):4289–96. doi: 10.1128/IAI.68.7.4289-4296.2000. PubMed PMID: 10858247; PubMed Central PMCID: PMC101748.

20. Copenhaver AM, Casson CN, Nguyen HT, Fung TC, Duda MM, Roy CR, et al. Alveolar macrophages and neutrophils are the primary reservoirs for *Legionella pneumophila* and mediate cytosolic surveillance of type IV secretion. Infect Immun. 2014;82(10):4325–36. Epub 20140804. doi: 10.1128/IAI.01891-14. PubMed PMID: 25092908; PubMed Central PMCID: PMC4187856.

21. Lettinga KD, Florquin S, Speelman P, van Ketel R, van der Poll T, Verbon A. Toll-like receptor 4 is not involved in host defense against pulmonary *Legionella pneumophila* infection in a mouse model. J Infect Dis. 2002;186(4):570–3. Epub 20020724. doi: 10.1086/341780. PubMed PMID: 12195388.

22. Mascarenhas DP, Zamboni DS. Innate immune responses and monocyte-derived phagocyte recruitment in protective immunity to pathogenic bacteria: insights from *Legionella pneumophila*. Curr Opin Microbiol. 2024;80:102495. Epub 20240621. doi: 10.1016/j.mib.2024.102495. PubMed PMID: 38908045.

23. Gonçalves AV, Margolis SR, Quirino GFS, Mascarenhas DPA, Rauch I, Nichols RD, et al. Gasdermin-D and Caspase-7 are the key Caspase-1/8 substrates downstream of the NAIP5/NLRC4 inflammasome required for restriction of *Legionella pneumophila*. PLoS Pathog. 2019;15(6):e1007886. Epub 20190628. doi: 10.1371/journal.ppat.1007886. PubMed PMID: 31251782; PubMed Central PMCID: PMC6622555.

24. Plumlee CR, Duffy, F. J., Gern, B. H., Delahaye, J. L., Cohen, S. B., Stoltzfus, C. R., Rustad, T. R., Hansen, S. G., Axthelm, M. K., Picker, L. J., Aitchison, J. D., Sherman, D. R., Ganusov, V. V., Gerner, M. Y., Zak, D. E., & Urdahl, K. B. Ultra-low Dose Aerosol Infection of Mice with *Mycobacterium tuberculosis* More Closely Models Human Tuberculosis. Cell host & microbe. 2021;29(1):68–82. PubMed Central PMCID: PMC7854984.

25. Skerrett SJ, Bagby GJ, Schmidt RA, Nelson S. Antibody-mediated depletion of tumor necrosis factor-alpha impairs pulmonary host defenses to *Legionella pneumophila*. J Infect Dis. 1997;176(4):1019–28. doi: 10.1086/516530. PubMed PMID: 9333161.

26. Barry KC, Fontana MF, Portman JL, Dugan AS, Vance RE. IL-1α signaling initiates the inflammatory response to virulent *Legionella pneumophila* in vivo. J Immunol. 2013;190(12):6329–39. Epub 20130517. doi: 10.4049/jimmunol.1300100. PubMed PMID: 23686480; PubMed Central PMCID: PMC3682686.

27. Blanchard DK, Klein TW, Friedman H, Stewart WE. Kinetics and characterization of interferon production by murine spleen cells stimulated with *Legionella pneumophila* antigens. Infect Immun. 1985;49(3):719–23. doi: 10.1128/iai.49.3.719-723.1985. PubMed PMID: 2411662; PubMed Central PMCID: PMC261255.

28. Martin CJ, Cadena AM, Leung VW, Lin PL, Maiello P, Hicks N, et al. Digitally Barcoding *Mycobacterium tuberculosis* Reveals In Vivo Infection Dynamics in the Macaque Model of Tuberculosis. mBio. 2017;8(3). Epub 20170509. doi: 10.1128/mBio.00312-17. PubMed PMID: 28487426; PubMed Central PMCID: PMC5424202.

29. Plumlee CR, Barrett HW, Shao DE, Lien KA, Cross LM, Cohen SB, et al. Assessing vaccine-mediated protection in an ultra-low dose *Mycobacterium tuberculosis* murine model. PLoS Pathog. 2023;19(11):e1011825. Epub 20231127. doi: 10.1371/journal.ppat.1011825. PubMed PMID: 38011264; PubMed Central PMCID: PMC10703413.

30. Beura LK, Hamilton SE, Bi K, Schenkel JM, Odumade OA, Casey KA, et al. Normalizing the environment recapitulates adult human immune traits in laboratory mice. Nature. 2016;532(7600):512–6. Epub 20160420. doi: 10.1038/nature17655. PubMed PMID: 27096360; PubMed Central PMCID: PMC4871315.

31. Huggins MA, Sjaastad FV, Pierson M, Kucaba TA, Swanson W, Staley C, et al. Microbial Exposure Enhances Immunity to Pathogens Recognized by TLR2 but Increases Susceptibility to Cytokine Storm through TLR4 Sensitization. Cell Rep. 2019;28(7):1729–43.e5. doi: 10.1016/j.celrep.2019.07.028. PubMed PMID: 31412243; PubMed Central PMCID: PMC6703181.

32. Labuda JC, Fong KD, McSorley SJ. Cohousing with Dirty Mice Increases the Frequency of Memory T Cells and Has Variable Effects on Intracellular Bacterial Infection. Immunohorizons. 2022;6(2):184–90. Epub 20220224. doi: 10.4049/immunohorizons.2100069. PubMed PMID: 35210292; PubMed Central PMCID: PMC9624231.

33. Flynn JL, Chan J, Triebold KJ, Dalton DK, Stewart TA, Bloom BR. An essential role for interferon gamma in resistance to *Mycobacterium tuberculosis* infection. J Exp Med. 1993;178(6):2249–54. doi: 10.1084/jem.178.6.2249. PubMed PMID: 7504064; PubMed Central PMCID: PMCPMC2191274.

34. Flynn JL, Goldstein MM, Chan J, Triebold KJ, Pfeffer K, Lowenstein CJ, et al. Tumor necrosis factor-alpha is required in the protective immune response against *Mycobacterium tuberculosis* in mice. Immunity. 1995;2(6):561–72. doi: 10.1016/1074-7613(95)90001-2. PubMed PMID: 7540941.

35. Ji DX, Witt KC, Kotov DI, Margolis SR, Louie A, Chevée V, et al. Role of the transcriptional regulator SP140 in resistance to bacterial infections via repression of type I interferons. Elife. 2021;10. Epub 20210621. doi: 10.7554/eLife.67290. PubMed PMID: 34151776; PubMed Central PMCID: PMC8248984.

36. Fremond CM, Yeremeev V, Nicolle DM, Jacobs M, Quesniaux VF, Ryffel B. Fatal *Mycobacterium tuberculosis* infection despite adaptive immune response in the absence of MyD88. J Clin Invest. 2004;114(12):1790–9. doi: 10.1172/JCI21027. PubMed PMID: 15599404; PubMed Central PMCID: PMC535064.

37. Watson RO, Bell SL, MacDuff DA, Kimmey JM, Diner EJ, Olivas J, et al. The Cytosolic Sensor cGAS Detects *Mycobacterium tuberculosis* DNA to Induce Type I Interferons and Activate Autophagy. Cell Host Microbe. 2015;17(6):811–9. Epub 20150602. doi: 10.1016/j.chom.2015.05.004. PubMed PMID: 26048136; PubMed Central PMCID: PMCPMC4466081.

38. Saini D, Hopkins GW, Seay SA, Chen CJ, Perley CC, Click EM, et al. Ultra-low dose of *Mycobacterium tuberculosis* aerosol creates partial infection in mice. Tuberculosis (Edinb). 2012;92(2):160–5. Epub 20111221. doi: 10.1016/j.tube.2011.11.007. PubMed PMID: 22197183; PubMed Central PMCID: PMC3288716.

39. Rao C, Benhabib H, Ensminger AW. Phylogenetic reconstruction of the *Legionella pneumophila* Philadelphia-1 laboratory strains through comparative genomics. PLoS One. 2013;8(5):e64129. Epub 20130522. doi: 10.1371/journal.pone.0064129. PubMed PMID: 23717549; PubMed Central PMCID: PMC3661481.

40. Molofsky AB, Byrne BG, Whitfield NN, Madigan CA, Fuse ET, Tateda K, et al. Cytosolic recognition of flagellin by mouse macrophages restricts *Legionella pneumophila* infection. J Exp Med. 2006;203(4):1093–104. Epub 20060410. doi: 10.1084/jem.20051659. PubMed PMID: 16606669; PubMed Central PMCID: PMC1584282.

41. Ren T, Zamboni DS, Roy CR, Dietrich WF, Vance RE. Flagellin-deficient *Legionella* mutants evade caspase-1- and Naip5-mediated macrophage immunity. PLoS Pathog. 2006;2(3):e18. Epub 20060317. doi: 10.1371/journal.ppat.0020018. PubMed PMID: 16552444; PubMed Central PMCID: PMC1401497.

42. Fontana MF, Banga S, Barry KC, Shen X, Tan Y, Luo ZQ, et al. Secreted bacterial effectors that inhibit host protein synthesis are critical for induction of the innate immune response to virulent *Legionella pneumophila*. PLoS Pathog. 2011;7(2):e1001289. Epub 20110217. doi: 10.1371/journal.ppat.1001289. PubMed PMID: 21390206; PubMed Central PMCID: PMCPMC3040669.

43. Stoeckius M, Hafemeister C, Stephenson W, Houck-Loomis B, Chattopadhyay PK, Swerdlow H, et al. Simultaneous epitope and transcriptome measurement in single cells. Nat Methods. 2017;14(9):865–8. Epub 20170731. doi: 10.1038/nmeth.4380. PubMed PMID: 28759029; PubMed Central PMCID: PMC5669064.

44. Büttner M, Ostner J, Müller CL, Theis FJ, Schubert B. scCODA is a Bayesian model for compositional single-cell data analysis. Nat Commun. 2021;12(1):6876. Epub 20211125. doi: 10.1038/s41467-021-27150-6. PubMed PMID: 34824236; PubMed Central PMCID: PMCPMC8616929.

45. Miller SA, Policastro RA, Sriramkumar S, Lai T, Huntington TD, Ladaika CA, et al. LSD1 and Aberrant DNA Methylation Mediate Persistence of Enteroendocrine Progenitors That Support. Cancer Res. 2021;81(14):3791–805. Epub 20210525. doi: 10.1158/0008-5472.CAN-20-3562. PubMed PMID: 34035083; PubMed Central PMCID: PMC8513805.

46. Liu D, Mai D, Jahn AN, Murray TA, Aitchison JD, Gern BH, et al. APOE protects against severe infection with *Mycobacterium tuberculosis* by restraining production of neutrophil extracellular traps. PLoS Pathog. 2025;21(6):e1013267. Epub 20250616. doi: 10.1371/journal.ppat.1013267. PubMed PMID: 40523023; PubMed Central PMCID: PMC12201663.

47. Lin PL, Ford CB, Coleman MT, Myers AJ, Gawande R, Ioerger T, et al. Sterilization of granulomas is common in active and latent tuberculosis despite within-host variability in bacterial killing. Nat Med. 2014;20(1):75–9. Epub 20131215. doi: 10.1038/nm.3412. PubMed PMID: 24336248; PubMed Central PMCID: PMC3947310.

48. MacMicking JD, Taylor GA, McKinney JD. Immune control of tuberculosis by IFN-gamma-inducible LRG-47. Science. 2003;302(5645):654–9. doi: 10.1126/science.1088063. PubMed PMID: 14576437.

49. Van Dis E, Fox DM, Morrison HM, Fines DM, Babirye JP, McCann LH, et al. IFN-γ-independent control of M. tuberculosis requires CD4 T cell-derived GM-CSF and activation of HIF-1α. PLoS Pathog. 2022;18(7):e1010721. Epub 20220725. doi: 10.1371/journal.ppat.1010721. PubMed PMID: 35877763; PubMed Central PMCID: PMC9352196.

50. Leal IS, Smedegård B, Andersen P, Appelberg R. Failure to induce enhanced protection against tuberculosis by increasing T-cell-dependent interferon-gamma generation. Immunology. 2001;104(2):157–61. doi: 10.1046/j.1365-2567.2001.01305.x. PubMed PMID: 11683955; PubMed Central PMCID: PMC1783293.

51. Sakai S, Kauffman KD, Sallin MA, Sharpe AH, Young HA, Ganusov VV, et al. CD4 T Cell-Derived IFN-γ Plays a Minimal Role in Control of Pulmonary *Mycobacterium tuberculosis* Infection and Must Be Actively Repressed by PD-1 to Prevent Lethal Disease. PLoS Pathog. 2016;12(5):e1005667. Epub 20160531. doi: 10.1371/journal.ppat.1005667. PubMed PMID: 27244558; PubMed Central PMCID: PMC4887085.

52. Srivastava S, Ernst JD. Cutting edge: Direct recognition of infected cells by CD4 T cells is required for control of intracellular *Mycobacterium tuberculosis* in vivo. J Immunol. 2013;191(3):1016–20. Epub 20130701. doi: 10.4049/jimmunol.1301236. PubMed PMID: 23817429; PubMed Central PMCID: PMC3725655.

53. Abel L, El-Baghdadi J, Bousfiha AA, Casanova JL, Schurr E. Human genetics of tuberculosis: a long and winding road. Philos Trans R Soc Lond B Biol Sci. 2014;369(1645):20130428. Epub 20140512. doi: 10.1098/rstb.2013.0428. PubMed PMID: 24821915; PubMed Central PMCID: PMC4024222.

54. Boisson-Dupuis S, Bustamante J. Mycobacterial diseases in patients with inborn errors of immunity. Curr Opin Immunol. 2021;72:262–71. Epub 20210724. doi: 10.1016/j.coi.2021.07.001. PubMed PMID: 34315005; PubMed Central PMCID: PMC9172628.

55. Azad AK, Sadee W, Schlesinger LS. Innate immune gene polymorphisms in tuberculosis. Infect Immun. 2012;80(10):3343–59. Epub 20120723. doi: 10.1128/IAI.00443-12. PubMed PMID: 22825450; PubMed Central PMCID: PMC3457569.

56. Setiabudiawan TP, Apriani L, Verrall AJ, Utami F, Schneider M, Indrati AR, et al. Immune correlates of early clearance of *Mycobacterium tuberculosis* among tuberculosis household contacts in Indonesia. Nat Commun. 2025;16(1):309. Epub 20250102. doi: 10.1038/s41467-024-55501-6. PubMed PMID: 39747050; PubMed Central PMCID: PMC11695729.

57. Lu LL, Smith MT, Yu KKQ, Luedemann C, Suscovich TJ, Grace PS, et al. IFN-γ-independent immune markers of *Mycobacterium tuberculosis* exposure. Nat Med. 2019;25(6):977–87. Epub 20190520. doi: 10.1038/s41591-019-0441-3. PubMed PMID: 31110348; PubMed Central PMCID: PMC6559862.

58. Dallmann-Sauer M, Fava VM, Malherbe ST, MacDonald CE, Orlova M, Kroon EE, et al. *Mycobacterium tuberculosis* resisters despite HIV exhibit activated T cells and macrophages in their pulmonary alveoli. J Clin Invest. 2025;135(7). Epub 20250121. doi: 10.1172/JCI188016. PubMed PMID: 39836471; PubMed Central PMCID: PMC11957701.

59. Li H, Wang XX, Wang B, Fu L, Liu G, Lu Y, et al. Latently and uninfected healthcare workers exposed to TB make protective antibodies against *Mycobacterium tuberculosis*. Proc Natl Acad Sci U S A. 2017;114(19):5023–8. Epub 20170424. doi: 10.1073/pnas.1611776114. PubMed PMID: 28438994; PubMed Central PMCID: PMC5441709.

60. Kotov DI, Lee OV, Fattinger SA, Langner CA, Guillen JV, Peters JM, et al. Early cellular mechanisms of type I interferon-driven susceptibility to tuberculosis. Cell. 2023;186(25):5536–53.e22. Epub 20231128. doi: 10.1016/j.cell.2023.11.002. PubMed PMID: 38029747; PubMed Central PMCID: PMC10757650.

61. Golovkine GR, Roberts AW, Morrison HM, Rivera-Lugo R, McCall RM, Nilsson H, et al. Autophagy restricts *Mycobacterium tuberculosis* during acute infection in mice. Nat Microbiol. 2023;8(5):819–32. Epub 20230410. doi: 10.1038/s41564-023-01354-6. PubMed PMID: 37037941; PubMed Central PMCID: PMCPMC11027733.

62. Green AM, Difazio R, Flynn JL. IFN-γ from CD4 T cells is essential for host survival and enhances CD8 T cell function during *Mycobacterium tuberculosis* infection. J Immunol. 2013;190(1):270–7. Epub 20121210. doi: 10.4049/jimmunol.1200061. PubMed PMID: 23233724; PubMed Central PMCID: PMCPMC3683563.

63. Fattinger SA, Chavez RA, Witt KC, Parisi B, Rodriguez JJ, Turcotte EA, et al. Strong sustained type I IFN signaling acts cell intrinsically to impair IFNγ responses and cause tuberculosis susceptibility. bioRxiv. 2026. Epub 20260122. doi: 10.64898/2026.01.19.700487. PubMed PMID: 41659471; PubMed Central PMCID: PMCPMC12879652.

64. Azuma M, Ito D, Yagita H, Okumura K, Phillips JH, Lanier LL, et al. B70 antigen is a second ligand for CTLA-4 and CD28. Nature. 1993;366(6450):76–9. doi: 10.1038/366076a0. PubMed PMID: 7694153.

65. Freeman GJ, Gribben JG, Boussiotis VA, Ng JW, Restivo VA, Lombard LA, et al. Cloning of B7-2: a CTLA-4 counter-receptor that costimulates human T cell proliferation. Science. 1993;262(5135):909–11. doi: 10.1126/science.7694363. PubMed PMID: 7694363.

66. Flores-Batista VC, Boechat N, Lago PM, Lazzarini LC, Pessanha LR, Almeida AS, et al. Low expression of antigen-presenting and costimulatory molecules by lung cells from tuberculosis patients. Braz J Med Biol Res. 2007;40(12):1671–9. Epub 20071029. doi: 10.1590/s0100-879×2006005000141. PubMed PMID: 17713660.

67. Vance RE. Tuberculosis as an unconventional interferonopathy. Curr Opin Immunol. 2025;92:102508. Epub 20241204. doi: 10.1016/j.coi.2024.102508. PubMed PMID: 39637776.

68. Moreira-Teixeira L, Mayer-Barber K, Sher A, O’Garra A. Type I interferons in tuberculosis: Foe and occasionally friend. J Exp Med. 2018;215(5):1273–85. Epub 20180417. doi: 10.1084/jem.20180325. PubMed PMID: 29666166; PubMed Central PMCID: PMCPMC5940272.

69. Lippmann J, Müller HC, Naujoks J, Tabeling C, Shin S, Witzenrath M, et al. Dissection of a type I interferon pathway in controlling bacterial intracellular infection in mice. Cell Microbiol. 2011;13(11):1668–82. Epub 20110824. doi: 10.1111/j.1462-5822.2011.01646.x. PubMed PMID: 21790939; PubMed Central PMCID: PMC3196383.

70. Stetson DB, Medzhitov R. Recognition of cytosolic DNA activates an IRF3-dependent innate immune response. Immunity. 2006;24(1):93–103. doi: 10.1016/j.immuni.2005.12.003. PubMed PMID: 16413926.

71. Medzhitov R, Schneider DS, Soares MP. Disease tolerance as a defense strategy. Science. 2012;335(6071):936–41. doi: 10.1126/science.1214935. PubMed PMID: 22363001; PubMed Central PMCID: PMCPMC3564547.

72. Ayres JS, Schneider DS. Tolerance of infections. Annu Rev Immunol. 2012;30:271–94. Epub 20120103. doi: 10.1146/annurev-immunol-020711-075030. PubMed PMID: 22224770.

73. Rhoades ER, Frank AA, Orme IM. Progression of chronic pulmonary tuberculosis in mice aerogenically infected with virulent *Mycobacterium tuberculosis*. Tuber Lung Dis. 1997;78(1):57–66. doi: 10.1016/s0962-8479(97)90016-2. PubMed PMID: 9666963.

74. Voskuil MI, Bartek IL, Visconti K, Schoolnik GK. The response of *Mycobacterium tuberculosis* to reactive oxygen and nitrogen species. Front Microbiol. 2011;2:105. Epub 20110513. doi: 10.3389/fmicb.2011.00105. PubMed PMID: 21734908; PubMed Central PMCID: PMC3119406.

75. Espich S, Berry S, Balakhmet A, Chang X, Thuong NTT, Dorajoo R, et al. Impaired aldehyde detoxification caused by a common human polymorphism promotes anti-bacterial immunity. bioRxiv. 2025. Epub Oct 18, 2025.

76. Thomas SM, Olive AJ. High Lethality of *Mycobacterium tuberculosis* Infection in Mice Lacking the Phagocyte Oxidase and Caspase1/11. Infect Immun. 2023;91(7):e0006023. Epub 20230614. doi: 10.1128/iai.00060-23. PubMed PMID: 37314361; PubMed Central PMCID: PMCPMC10353354.

77. Darwin KH. *Mycobacterium tuberculosis*: the honey badger of pathogens. EMBO Rep. 2021;22(9):e53619. Epub 20210728. doi: 10.15252/embr.202153619. PubMed PMID: 34322986; PubMed Central PMCID: PMC8419694.

78. Gordon C. The crazy nastyass honeybadger. Youtube.com 2011. p. https://www.youtube.com/watch?v=4r7wHMg5Yjg.

79. Mampel J, Spirig T, Weber SS, Haagensen JA, Molin S, Hilbi H. Planktonic replication is essential for biofilm formation by *Legionella pneumophila* in a complex medium under static and dynamic flow conditions. Appl Environ Microbiol. 2006;72(4):2885–95. doi: 10.1128/AEM.72.4.2885-2895.2006. PubMed PMID: 16597995; PubMed Central PMCID: PMC1448985.

80. Anaya-Sanchez A, Feng Y, Berude JC, Portnoy DA. Detoxification of methylglyoxal by the glyoxalase system is required for glutathione availability and virulence activation in *Listeria monocytogenes*. PLoS Pathog. 2021;17(8):e1009819. Epub 20210818. doi: 10.1371/journal.ppat.1009819. PubMed PMID: 34407151; PubMed Central PMCID: PMCPMC8372916.

81. Ashburner M, Ball CA, Blake JA, Botstein D, Butler H, Cherry JM, et al. Gene ontology: tool for the unification of biology. The Gene Ontology Consortium. Nat Genet. 2000;25(1):25–9. doi: 10.1038/75556. PubMed PMID: 10802651; PubMed Central PMCID: PMCPMC3037419.

82. Consortium GO. The Gene Ontology knowledgebase in 2026. Nucleic Acids Res. 2026;54(D1):D1779–D92. doi: 10.1093/nar/gkaf1292. PubMed PMID: 41413728; PubMed Central PMCID: PMCPMC12807639.

83. Xu S, Hu E, Cai Y, Xie Z, Luo X, Zhan L, et al. Using clusterProfiler to characterize multiomics data. Nat Protoc. 2024;19(11):3292–320. Epub 20240717. doi: 10.1038/s41596-024-01020-z. PubMed PMID: 39019974.

